# Orthosteric Molecular Glue Inhibits COP9 Signalosome with Substrate-Dependent Potency

**DOI:** 10.1101/2025.11.26.690573

**Authors:** Huigang Shi, Xiaorong Wang, Clinton Yu, Haibin Mao, Fenglong Jiao, Merav Braitbard, Ben Shor, Zhongsheng Zhang, Thomas R. Hinds, Shiyun Cao, Erkang Fan, Dina Schneidman-Duhovny, Lan Huang, Ning Zheng

**Affiliations:** Department of Pharmacology, Box 357280, University of Washington, Seattle, WA 98195, USA; Department of Physiology and Biophysics, University of California, Irvine, CA 92697, USA; Howard Hughes Medical Institute, University of Washington, Seattle, WA 98195, USA; The Rachel and Selim Benin School of Computer Science and Engineering, The Hebrew University of Jerusalem, Jerusalem, Israel; Department of Biochemistry, University of Washington, Seattle, WA 98175, USA

## Abstract

Orthosteric inhibitors block enzyme active sites and prevent substrates from binding. Enhancing their specificity through substrate dependence seems inherently unlikely, as their mechanism hinges on direct competition rather than selective recognition. Here, we show that a molecular glue mechanism unexpectedly imparts substrate-dependent potency to CSN5i-3, an orthosteric inhibitor of the COP9 signalosome (CSN). We first confirm that CSN5i-3 inhibits CSN, which catalyzes NEDD8 deconjugation from the cullin-RING ubiquitin ligases (CRLs), by occupying the active site of its catalytic subunit, CSN5, and directly competing with the iso-peptide bond substrate. Curiously, the orthosteric inhibitor binds free CSN with only micromolar affinity, yet achieves nanomolar potency in blocking its deneddylase activity. Cryo-EM structures of the enzyme-substrate-inhibitor complex reveal that active site-engaged CSN5i-3 occludes the substrate iso-peptide linkage while simultaneously extending an NEDD8-binding exosite of CSN5, acting as a molecular glue to cement the NEDD8-CSN5 interaction. The cooperativity of this tri-molecular CSN5i-3-NEDD8-CSN5 assembly, in turn, sequesters CSN5i-3 at its binding site, conferring high potency to the orthosteric inhibitor despite its low affinity for the free enzyme. Together, our findings highlight the modest affinity requirements of molecule glues for individual target proteins and establish “orthosteric molecular glue inhibitors” as a new class of substrate-dependent enzyme antagonists.

## INTRODUCTION

Enzymatic inhibitors are indispensable tools for dissecting biological pathways and developing therapeutic interventions^1^. They are broadly categorized by their binding sites and mechanisms of action. Among these, orthosteric inhibitors, which bind to the catalytic site and directly compete with substrates, have been extensively explored due to their predictable structure-activity relationships. However, such inhibitors are typically substrate-agnostic, as their mechanism relies solely on blocking the active site. In contrast, substrate-dependent inhibitors, which achieve selectivity by engaging allosteric sites or exosites, can modulate enzyme activity in a substrate-specific manner^2^. Yet, their design remains challenging due to the complex structural and dynamic determinants governing these interactions. Ideally, combining the tractability of orthosteric inhibitors with the precision of substrate-dependent modulation would offer a powerful strategy—but whether such a hybrid approach is feasible has remained unclear.

The COP9 signalosome (CSN) is a multi-subunit protein complex evolutionarily conserved across all eukaryotic species^3,4^. It plays a crucial role in regulating almost every aspect of cellular functions by modulating the cullin-RING ubiquitin ligases (CRLs), the major components of the ubiquitin-proteasome system. In plants, CSN is essential for suppressing photomorphogenesis in the dark^5^. In humans, CSN controls the E3 activity of hundreds of CRLs in diverse cellular pathways and is required for targeted protein degradation induced by emerging protein degraders, including PROTACs and molecular glues^6–12^. As the largest family of E3s in eukaryotic cells, the proper assembly and the ubiquitin ligase activity of the CRL E3s hinges on the dynamic modification of the cullin scaffolds by a ubiquitin-like protein, NEDD8 (N8), also known as neddylation^13^. While cullin neddylation has been reported to enhance the E3 activity of CRLs, CSN-mediated cullin deneddylation is thought to protect the CRL SRs from auto-ubiquitination and promote their exchange on the cullin scaffolds through an adaptive CRL assembly cycle^14,15^.

CSN comprises eight core subunits, CSN1–CSN8 (**Fig. 1a**). With a conserved JAB1 MPN domain metalloenzyme (JAMM) motif, CSN5 functions as the sole catalytic subunit of the deneddylase complex^16^. The crystal structure of the CSN holoenzyme revealed that CSN5 and CSN6 form a heterodimer, which is affixed to the other six subunits that oligomerize via their winged-helix subdomains and C-terminal bundle helices^17^. Interestingly, the active site of CSN5 is occluded by its insertion-1 (Ins-1) loop in the CSN holoenzyme structure, suggesting that the iso-peptidase complex might be in an “inactive” state. Multiple subsequent cryo-EM studies have since captured the overall architecture of CSN in complexes with four different N8∼CRLs^18–22^. These structures revealed a catalytic hemisphere of CSN formed among CSN2, CSN4, CSN5, and CSN6 that undergoes major conformational changes upon substrate engagement (**Fig. 1b**). All these structures, however, suffer from a limited resolution, especially within the catalytic hemisphere. How the NEDD8∼cullin iso-peptide linkage is recognized by the catalytic site of CSN in its “pre-catalytic state” remains to be elucidated.

**Fig. 1.**
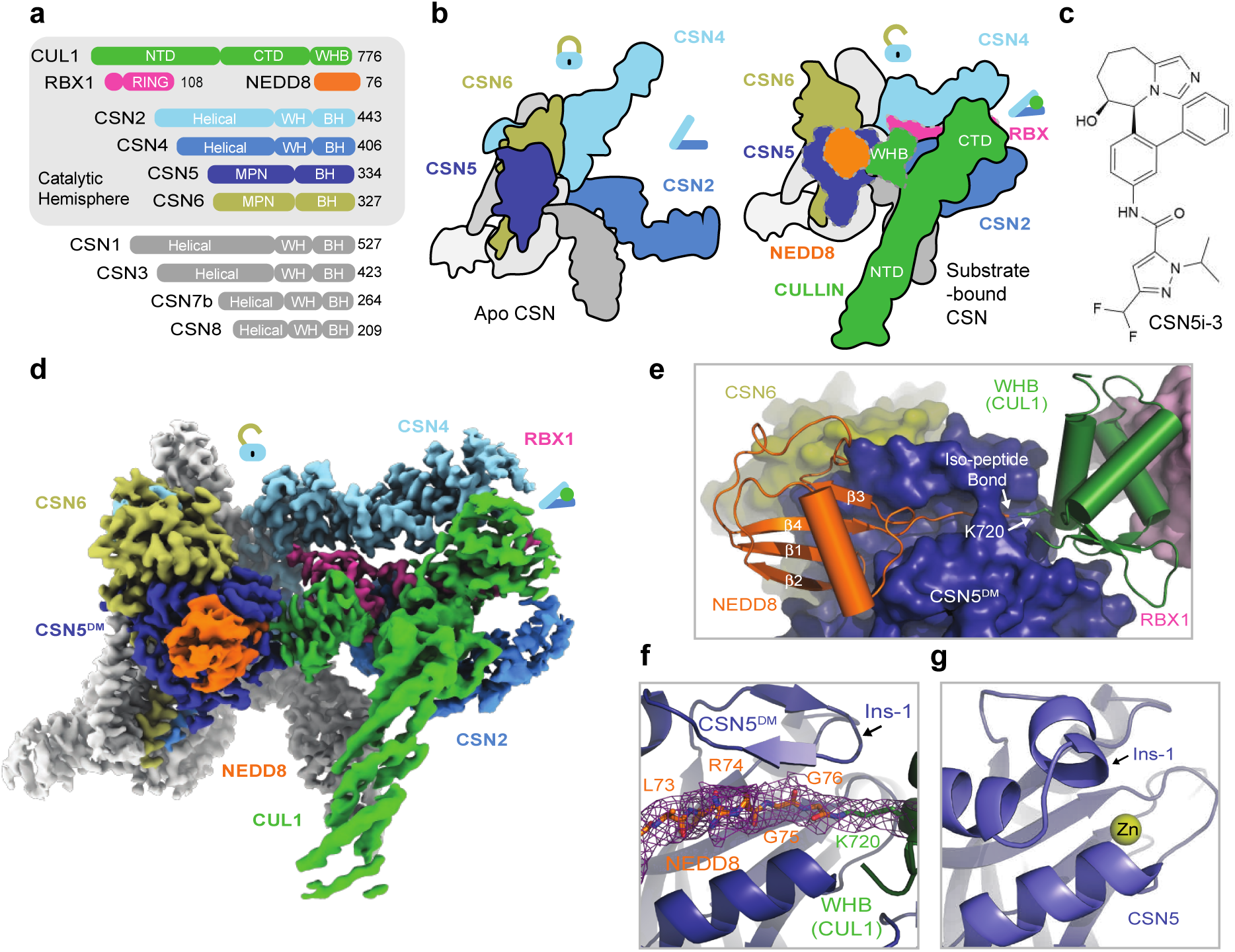
The pre-catalytic state of CSN in complex with N8∼CRL1. **a.** Domain composition of CSN subunits, NEDD8, and the CUL1-RBX1 E3 scaffold. **b.** Schematic drawings of CSN in its apo and N8∼cullin-RING-bound forms with the CSN subunits in the catalytic hemisphere and the N8∼CRL substrate colored. Major conformational changes in CSN upon substrate engagement are indicated by lock and clamp icons. **c.** The chemical structure of CSN5i-3. **d.** Cryo-EM map of the CSN^DM^-N8∼CRL1 complex. **e.** The three-molecular interface among CSN5^DM^ (surface in dark blue), NEDD8 (cartoon in orange), and the CUL1-WHB domain (carton in green). The iso-peptide bond formed between the NEDD8 C-terminal carboxyl group and the lysine residue, Lys120, of CUL1 is indicated. **f.** Recognition of the iso-peptide bond formed between NEDD8 and Lys720 of the CUL1-WHB domain by CSN5^DM^ with its reshaped Ins-1 loop. **g.** A close-up view of the catalytic site in isolated CSN5 occupied by its Ins-1 loop. PDB:4D10.

With multiple subunits overexpressed in many human cancers, CSN has been widely recognized as a potential target for anti-cancer therapies. Indeed, CSN5i-3 has recently been developed as a small molecule CSN5 inhibitor that can potently block cullin deneddylation by CSN (**Fig. 1c**)^23^. Although the compound has been shown to directly target the active site of CSN5, it has recently been reported to inhibit the deneddylase complex via a surprisingly uncompetitive mechanism^24^. Unlike a noncompetitive inhibitor, an uncompetitive (also known as anti-competitive) inhibitor is expected to bind only to the enzyme-substrate complex, not to the free enzyme^25^. In this study, we discover that CSN5i-3 unexpectedly acts as a molecular glue, which not only stabilizes the CSN-N8∼CRL complexes but also gains its high potency in a substrate-dependent manner. This unusual mechanism of action leads us to establish the concept of “orthosteric molecular glue inhibitors”.

## RESULTS

### Pre-catalytic state structure of CSN^DM^-N8∼CRL1

As a metalloprotease, CSN5 features a catalytic Zn^2+^ ion at the end of a hydrophobic cleft, which has been predicted to recognize the C-terminal tail of NEDD8 conjugated to cullins^18^. A crystal structure of isolated CSN5 bound to CSN5i-3 revealed that the compound directly coordinates the Zn^2+^ ion and occupies the enzyme active site as an orthosteric inhibitor^23^. Before reconciling its orthosteric nature and its reported uncompetitive mechanism, we first set to experimentally determine how the N8∼CRL1 iso-peptide linkage is physically engaged with the CSN5 catalytic cleft in the absence of the compound. The catalytic zinc ion in CSN5 is coordinated by an aspartate (Asp151) and two histidine residues (His138 and His140), which are joined by an upstream water-activating glutamate residue (Glu76)^16,17,26^. Using a catalytically impaired CSN mutant harboring a CSN5 H138A mutation, previous studies have not been able to resolve the binding mode of the N8∼cullin conjugates at the CSN5 catalytic site. In an enzymatic assay with N8∼CRL1 as a substrate, we found that the CSN^CSN5-H138A^ mutant, as well as several other documented CSN mutants, still retains a detectable iso-peptidase activity (**Extended Fig. 1a**). By contrast, a CSN5 E76A/D151N double mutation completely abolishes the catalytic activity of CSN towards N8∼CRL1. Leveraging this CSN mutant (hereafter referred to as CSN^DM^), we determined the cryo-EM structure of a CSN^DM^-N8∼CRL1 complex at a resolution of 3.2 Å (**Fig. 1d, Extended Data Fig. 1b, Extended Data Table 1**).

Distinct from all previously reported CSN-N8∼CRL structures, the catalytic hemisphere of the CSN^DM^-N8∼CRL1 complex is clearly resolved in the 3D reconstruction map with the substrate iso-peptide linkage stably trapped at the CSN5 catalytic site (**Fig. 1e**). The resulting structural model confirms the previously described global architectural changes in CSN upon binding N8∼CRLs, which are highlighted by the pivotal movement of the CSN5-CSN6 dimer induced by the association of CUL1-CTD-RBX1 with CSN2 and CSN4. In addition, the complex structure reveals a cascade of protein-protein interactions around the catalytic center, involving NEDD8, CSN5, CUL1-WHB, and the RBX1 RING domain. Importantly, the resolution of the structure is high enough to resolve the complete NEDD8 C-terminal tail conjugated to the side chain of the CUL1-WHB Lys720 residue, which together span the entire CSN5 catalytic cleft. In comparison to the apo form of CSN, the central region of the CSN5 Ins-1 loop is dislodged from the catalytic cleft and adopts a β-hairpin structure, forming a three-stranded anti-parallel β-sheet with NEDD8 C-terminal tail (**Fig. 1f, g**). Such a structural arrangement positions the N8∼CRL1 iso-peptide bond right above the catalytic zinc ion binding site, representing the pre-catalytic state of the enzyme-substrate complex.

### Pre-catalytic state protein-protein interactions

Similar to all previously reported CSN-N8∼CRL structures, the polypeptide sequence connecting the CUL1 WHB domain to the CTD has no visible density and is presumably flexible in structure. The CUL1 WHB domain, nevertheless, is well resolved in the cryo-EM map, forming an extensive interface with CSN5 centered around the neddylation site, CUL1-Lys720 (**Fig. 2a, b**). Part of this interface is made between CUL1-WHB and the CSN5 MPN core domain, where the aliphatic side chain of CUL1-Lys720 is buttressed by two aromatic residues of CSN5, Tyr143 and Trp146. These two CSN5 residues, at the same time, interact with Met721 and Lys723 of CUL1-WHB via hydrophobic and π-cation interactions, respectively. Two additional CSN5 residues at the tip of the Ins-1 loop β-hairpin, Glu104 and Thr105, further strengthen the interface by interacting with Arg717 of CUL1-WHB via a salt bridge and van der Waal packing (**Fig. 2a**). Upon engaging N8∼CRL1, the rotation movement of the CSN5-CSN6 dimer pivots the CSN5 Ins-2 loop away from the helical bundle. Adopting a new conformation, this second CSN5 insertion loop joins Trp146 to wrap around the aliphatic side chain of CUL1-Lys723 (**Fig. 2b**). Two of its hydrophobic residues, Ile211 and Val216, further augment the interface via hydrophobic packing.

**Fig. 2.**
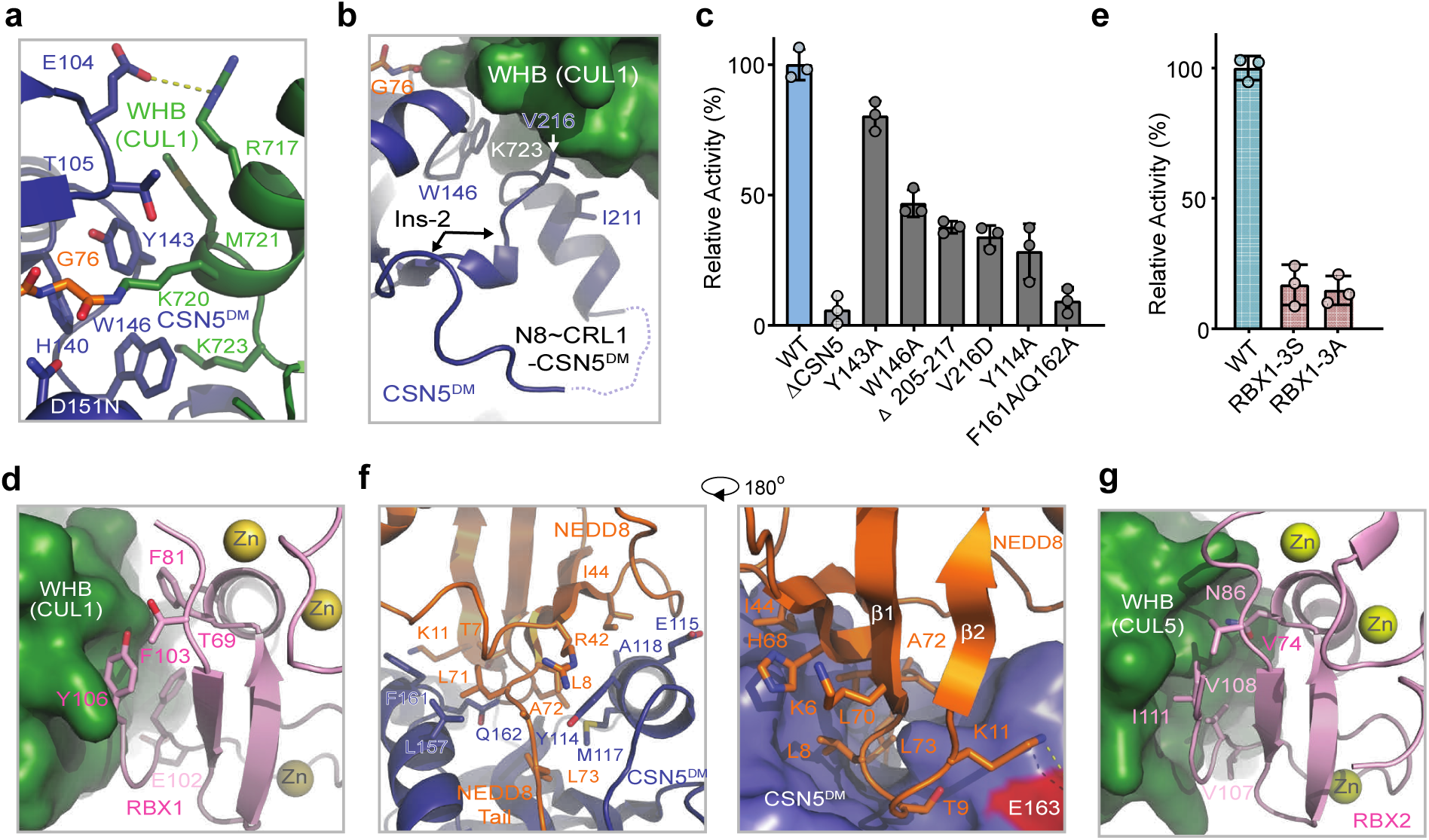
Protein interactions stabilizing the pre-catalytic state of CSN-N8∼CRL1. **a.** A close-up view of the interface between CSN5^DM^ and the CUL1-WHB domain with the substrate’s iso-peptide bond housed at the catalytic site. **b.** A close-up view of the CSN5 Ins-2 loop in the CSN^DM^-N8∼CRL1 complex. **c.** A comparison of the enzymatic activities of CSN containing WT and mutant CSN5. **d.** A close-up view of the WHB-RBX1 interface. **e.** A comparison of the enzymatic activities of CSN towards N8∼CRL1 featuring WT and mutant RBX1 with Phe81, Phe103, and Y106 mutated to serine or alanine. **f.** Two close-up views of the exosite interface between NEDD8 (orange) and CSN5^DM^ (slate). **g.** A close-up view of the interface between CUL5-WHB and RBX2-RING.

To test the functional importance of this CSN5-CUL1-WHB interface, we measured the enzymatic activity of CSN with the CSN5 Tyr143 and Trp146 residues individually mutated to alanine. Both mutants were catalytically compromised, albeit to different degrees (**Fig. 2c**). Similarly, an internal truncation of the CSN5 Ins-2 loop (Δ205-217), markedly attenuated the activity of the iso-peptidase complex. In fact, converting Val216, a hydrophobic residue in the Ins-2 loop, to a charged amino acid was sufficient to achieve the same effect (**Fig. 2c**). These results suggest that the CSN5-CUL1-WHB interaction is functionally coupled to the engagement of the iso-peptide linkage at the CSN5 catalytic cleft.

By securing the CUL1 WHB domain against CSN5, the active site-engaged iso-peptide linkage also stabilizes the packing of CUL1-WHB against the RBX1 RING domain at the opposite side (**Fig. 2d**). Characterized by a mixture of hydrophobic and polar interactions, this CUL1-WHB-RBX1-RING interface is mediated by a cluster of three aromatic residues, Phe81, Phe103, and Tyr106, which are strictly conserved in RBX1 orthologs (**Extended Data Fig. 1c**). Mutation of these residues to either serine or alanine effectively abolished the enzymatic activity of CSN without impacting N8∼CRL1 binding (**Fig. 2e, Extended Data Fig. 1d–f**). The optimal enzymatic activity of CSN, therefore, requires a continuum of intermolecular interfaces extending from the CSN5 active site to the RBX1 RING domain, which itself is locked by CSN2 and CSN4 as shown by previous studies^18^.

### NEDD8-binding exosite on CSN5

Besides the two separate interfaces made by the CUL1-WHB domain with CSN5 and the RBX1 RING domain, immobilization of the iso-peptide conjugate at the CSN5 active site also stabilizes the NEDD8 globular domain, which is anchored to an exosite of CSN5 next to the catalytic cleft. Immediately preceding the six amino acids NEDD8 C-terminal tail (Leu71–Gly76), the β1-β2 loop of the NEDD8 globular domain is wedged into a surface pocket of CSN5 adjacent to its catalytic cleft (**Fig. 2f**). On the two sides of this loop are two overall hydrophobic patches, including one centered around Ile44, that mediate the recognition of NEDD8 by CSN5. The tip of the loop is further locked in place by a network of hydrogen bonds and salt bridges. Interestingly, a missense mutation of a CSN5 glutamate residue at this interface (E163 in human CSN5) causes defects in photoreceptor neuron projections in *Drosophila*^27^. Despite anchoring at a different angle, the binding mode of the NEDD8 globular domain to CSN5 resembles the recognition of the distal ubiquitin by AMSH-LP, a JAMM-type zinc-dependent deubiquitinase (DUB) capable of cleaving Lys63-linked polyubiquitin chain (**Extended Data Fig. 1g**)^26^. Intriguingly, all residues within the two modifiers that are perceived by their cognate isopeptidases are conserved between NEDD8 and ubiquitin, even though the majority of the modifier-binding amino acids are different in the two enzymes (**Extended Data Fig. 1h**). This exosite interface between NEDD8 and CSN5, therefore, is unlikely the main determinant of the substrate specificity of CSN. In agreement, CSN has been previously shown to be able to hydrolyze the artificial ubiquitin-rhodamine substrate^17^. Although this NEDD8-CSN5 exosite interface cannot differentiate NEDD8 from ubiquitin, it is crucial for cullin deneddylation. Mutations of the primary NEDD8-contacting residues in CSN5, i.e., Tyr114, Phe161, and Gln162, impair or even abrogate the enzymatic activity of CSN (**Fig. 2c**). The productive engagement between the deneddylase complex and its substrate, therefore, entails a network of highly coupled protein-protein interactions traversing from the NEDD8-binding exosite on CSN5 to CSN2/4 via the iso-peptide linkage, the CUL1 WHB domain, and the RBX1 RING domain.

### Pre-catalytic states of all CSN-N8∼CRLs

To confirm that the pre-catalytic architecture of CSN-N8∼CRL1 is applicable to other CRLs, we obtained the cryo-EM structures of CSN^DM^ in complexes with NEDD8-modified CUL2-RBX1, CUL3-RBX1, CUL4A-RBX1, and CUL5-RBX2. Apart from the CSN^DM^-N8∼CRL3 structure, which was resolved at a 3.9 Å resolution, all other structures were determined with a high enough resolution (3.0 – 3.5 Å) to reveal the detailed interfaces within the catalytic hemisphere (**Extended Data Fig. 2a–d, Extended Data Table 1**). As expected, these structures share the same topology as the CSN^DM^-N8∼CRL1 complex, which is highlighted by the same chain of interactions that bridges NEDD8 and CSN2/4. Interestingly, the three aromatic residues of RBX1 interfacing with the CUL-1WHB domain are replaced by three non-aromatic hydrophobic residues in RBX2 (**Fig. 2g**). The same RING-WHB binding pose is, nevertheless, retained in the CSN^DM^-N8∼CRL5 complex. A superposition analysis reveals that the WHB domains of CUL1 and CUL5 are positioned slightly differently relative to their flanking CSN5 and RING partners (**Extended Fig. 1i**). Such a topological variation is accommodated by the differential pivotal angles of the CSN5-CSN6 dimer related to the rest of CSN. The iso-peptidase complex, therefore, harnesses both its intrinsic plasticity and synchronized protein-protein interactions to catalyze the deneddylation reaction across all cullin proteins.

### CSN5i-3 as a hybrid orthosteric uncompetitive inhibitor

After revealing how the substrate and its iso-peptide interact with the CSN active site, we next determined the CSN5i-3-CSN complex structure by cryo-EM at a resolution of 3.3 Å to verify the binding mode of the inhibitor in the context of the intact deneddylase complex (**Fig. 3a Extended Data Fig. 3a, Extended Data Table 1**). The structure of CSN bound to CSN5i-3 is nearly identical to its apo form, except that the inhibitory compound is sufficient to displace the Ins-1 loop and gain access to the active site without the engagement of a neddylated cullin-RING substrate (**Fig. 3b**). CSN5i-3 adopts the same binding pose on CSN5 within the CSN complex as it does on isolated CSN5. When superimposed with the CSN^DM^-N8∼CLR1 structure, CSN5i-3 would severely clash with the NEDD8 C-terminal tail, the iso-peptide bond, and the CUL1-Lys720 side chain, confirming its orthosteric nature (**Fig. 3c**).

**Fig. 3.**
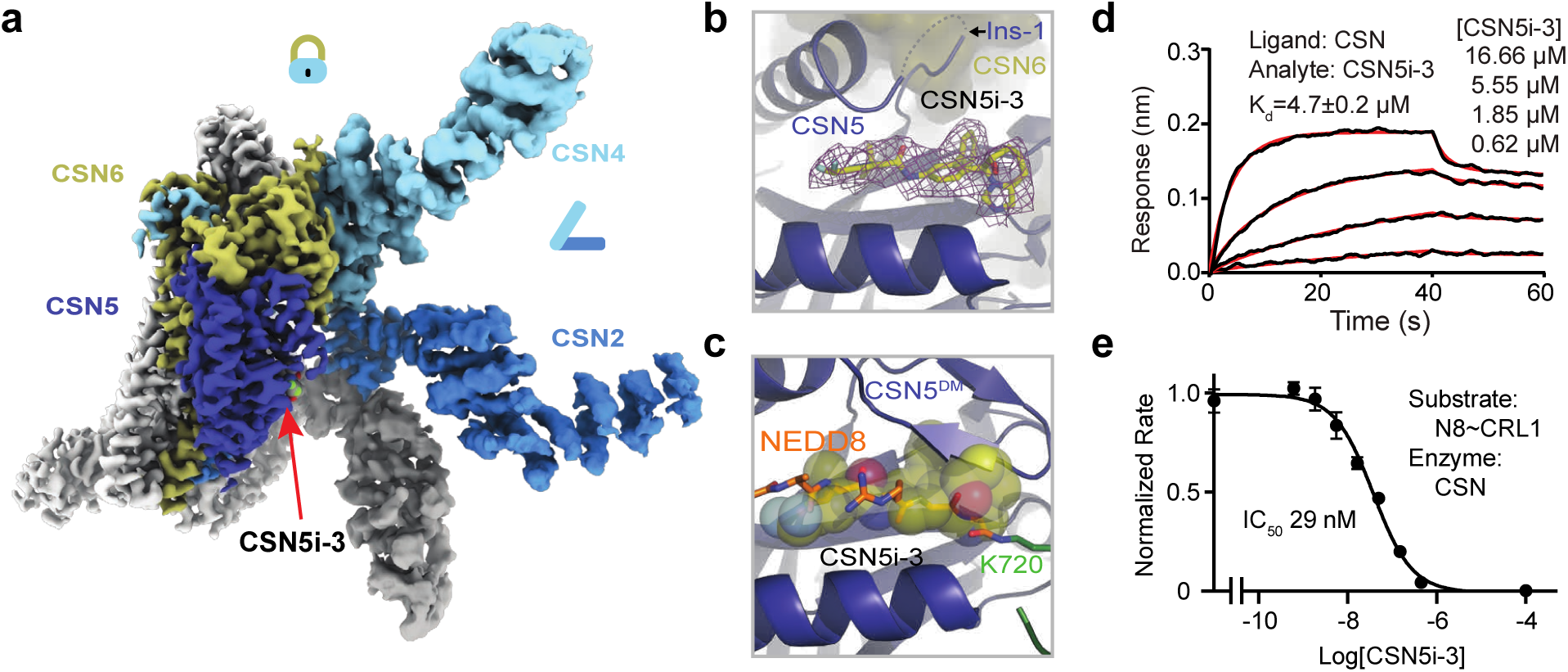
CSN5i-3 as a hybrid orthosteric uncompetitive inhibitor of CSN. **a.** Cryo-EM map of CSN5i-3-bound CSN with subunits in the catalytic hemisphere colored and the inhibitor shown in spheres. **b.** A close-up view of CSN5 catalytic cleft occupied by CSN5i-3, the density of which is shown in purple mesh. **c.** Steric hindrance imposed by CSN5i-3 for the engagement of the N8∼CRL1 iso-peptide linkage at the CSN5 active site. **d.** The binding affinity of CSN5i-3 to CSN measured by BLI. **e.** The potency of CSN5i-3 in inhibiting the iso-peptidase activity of CSN towards N8∼CRL1 in vitro.

To further characterize the inhibitor, we used biolayer interferometry (BLI) and measured the affinity of CSN5i-3 to a purified CSN5-CSN6 heterodimer. Strikingly, the dissociation constant, *K_D_,* of the inhibitor-CSN5-CSN6 interaction was determined to be ∼4 μM (**Extended Data Fig. 3b**), which differs from the originally reported potency of CSN5i-3 (∼5.8 nM) by three orders of magnitude^23^. This unexpected moderate affinity of the inhibitor was further verified by isothermal titration calorimetry (ITC) (**Extended Data Fig. 3c**). Consistent with its identical binding mode on isolated and CSN-embedded CSN5, the inhibitor displayed a similar affinity (∼4.7 μM) towards the free eight-subunits CSN complex (**Fig. 3d**). Using N8∼CRL1 as a substrate, we further confirmed that CSN5i-3 inhibited the CSN complex for deconjugating N8∼CRL1 with a low nM IC_50_ (29 nM) (**Fig. 3e**). CSN5i-3, therefore, acts as a noncanonical uncompetitive inhibitor that preferentially binds to the enzyme-substrate complex rather than to the free enzyme. Based on the established model of CSN activation, the most parsimonious explanation would be substrate-induced remodeling of the auto-inhibitory Ins-1 loop, which might facilitate CSN5i-3 binding to the CSN5 active site.

### Structure of CSN5i-3-bound CSN-N8∼CRL1

To decipher the precise uncompetitive mechanism of CSN5i-3, we assembled and determined the cryo-EM structure of inhibitor-bound CSN in complex with N8∼CRL1 at a resolution of 3.0 Å (**Fig. 4a**, **Extended Data Fig. 3d**, **Extended Data Table 1**). As expected, when bound to the N8∼CRL1 complex, CSN5i-3 blocks the CSN5 catalytic cleft and pushes the substrate iso-peptide conjugate out of the active site. Although such an action of the orthosteric inhibitor is anticipated to disrupt the protein interaction network that stabilizes the pre-catalytic state of the enzyme-substrate assembly, the CSN-N8∼CLR1-CSN5i-3 complex structure reveals a well-organized catalytic hemisphere with clearly resolved densities for NEDD8, CUL1-WHB, and RBX1-RING. While preventing the substrate iso-peptide linkage from binding the CSN5 active site, the orthosteric inhibitor appears to make direct contacts with NEDD8 and CUL1-WHB and maintain the overall topology of the pre-catalytic assembly (**Fig. 4b-d**).

**Fig. 4.**
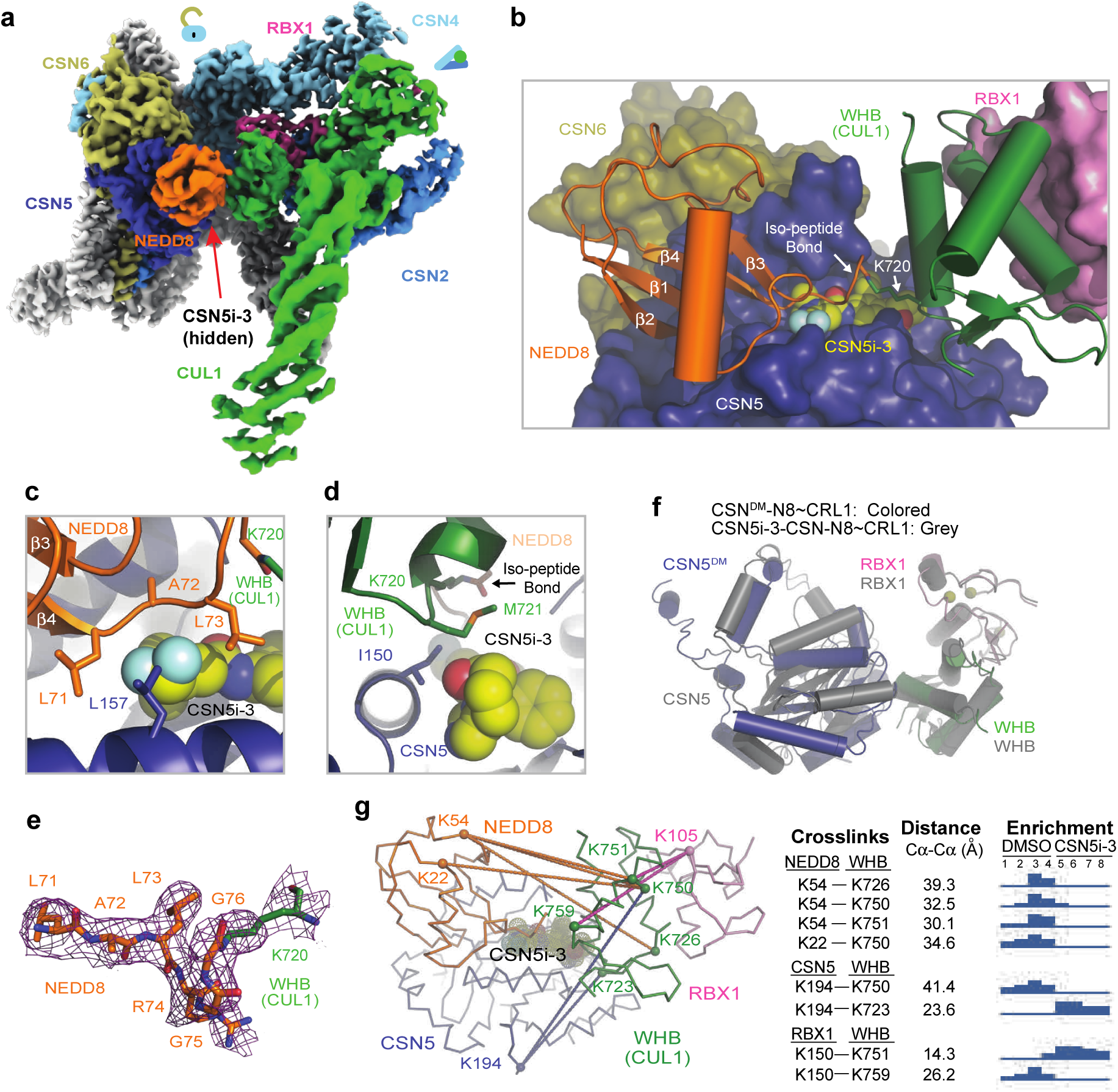
Stabilization of CSN-N8∼CRL1 pre-catalytic state by CSN5i-3. **a.** Cryo-EM map of the CSN5i-3-CSN-N8∼CRL1 complex with the substrate and CSN subunits in the catalytic hemisphere colored. **b.** The four-molecular interface among CSN5i-3 (sphere), CSN5 (surface in dark blue), NEDD8 (cartoon in orange), and the CUL1-WHB domain (carton in green). The iso-peptide bond formed between the NEDD8 C-terminal carboxyl group and the lysine residue of CUL1 is indicated. **c.** A close-up view of the interface between CSN5i-3 and the C-terminal tail of NEDD8. **d.** A close-up view of the interface among CSN5i-3, CSN5, and the CUL1-WHB domain. **e.** The cryo-EM density of the iso-peptide bond flanked by the NEDD8 C-terminal tail and the side chain of the neddylation site lysine residue in CUL1-WHB domain. **f.** Positional difference of CSN5 relative to the CUL1-WHB domain anchored to the RBX1 RING domain in the structures of CSN^DM^ and CSN5i-3-bound CSN in complex with N8∼CRL1. **g.** Select crosslinks among NEDD8, CSN5, CUL1-WHB, and RBX1-RING mapped on the CSN5i-3-CSN-N8∼CRL1 cryo-EM structure along with the corresponding Cα-Cα distances and the enrichment profile detected by quantitative TMT mass spectrometry.

In complex with CSN-N8∼CRL1, the elongated CSN5i-3 compound adopts the same binding pose as observed in the free form of CSN, spanning the whole catalytic cleft and burying the catalytic zinc ion underneath (**Extended Data Fig. 3e**). At one end of the inhibitor, the difluoromethyl pyrazole moiety of CSN5i-3 packs closely against the N-terminal half of the NEDD8 C-terminal tail, which consists of three residues, Leu71, Ala72, and Leu73 (**Fig. 4c**). The difluoromethyl group, in particular, is sandwiched between the two leucine residues of NEDD8 and is further shielded by Leu157 of CSN5. Together, they nucleate a hydrophobic core and extend the adjacent NEDD8-CSN5 exosite interface (**Fig. 4b, Extended Data Fig. 3f**).

At the opposite end of the catalytic cleft, the otherwise solvent exposed azepine group of the inhibitor is engaged with Met721 from the CUL1 WHB domain, which is next to the neddylation site, Lys720. In conjunction with the nearby Ile150 residue in CSN5, they weld a separate tri-molecular hydrophobic interface (**Fig. 4d**). While being excluded from the catalytic cleft by CSN5i-3, the linkage between the C-terminal di-glycine motif of NEDD8 and the side chain of CUL1 Lys720 is clearly visible in the cryo-EM map, overarching the inhibitory compound (**Fig. 4b, 4e**). In comparison to the CSN^DM^-N8∼CRL1 structure, the CUL1 WHB domain maintains its interface with the RBX1 RING domain, while packing against CSN5 at a slightly different angle (**Fig. 4f**).

To verify the topology of the catalytic hemisphere of the CSN-N8∼CRL1 complex stabilized by CSN5i-3, we employed the QMIX (quantitation of multiplexed, isobaric-labeled cross (X)-linked peptides) strategy for multiplexed quantitative cross-linking mass spectrometry (XL-MS) analysis of CSN mixed with N8∼CRL1 in the presence and absence of the inhibitor^28^. An MS-cleavable lysine-reactive cross-linker, DSSO, was coupled with isobaric tandem mass tag (TMT) reagents for quantitative profiling of protein-protein interactions with residue specific resolution^29^. As expected, CSN5i-3 treatment enriched numerous inter-molecular XLs between CUL1 and NEDD8 likely by blocking the deneddylation reaction. A few crosslinks between the CUL1-WHB domain and NEDD8, nevertheless, were clearly suppressed by the inhibitor, consistent with their spatial separation as observed in the cryo-EM structure (**Fig. 4g, Extended Data Table 2**). Remarkably, CSN5i-3 elicited opposite effects on the crosslinks of a lysine residue in RBX1-RING as well as in CSN5 to two lysine residues in the CUL1-WHB domain. In both cases, the opposite effects can be explained by the maximal inter-lysine distance (∼ 30 Å) allowed for efficient XL (**Fig. 4g**). Our XL-MS data, therefore, fully supports the spatial architecture of the enzyme-substrate complex resolved by cryo-EM.

### CSN5i-3 as a molecular glue

Despite acting as an orthosteric inhibitor, CSN-bound CSN5i-3 retains the NEDD8-CUL1-WHB conjugate, which appears to block its exit path and trap it in the catalytic cleft. The interfacial nature of the compound hint at a cooperative binding mechanism characteristic of a molecular glue^30^. Specifically, the close proximity between CSN5i-3 and the NEDD8-CSN5 exosite interface strongly suggests that the iso-peptidase inhibitor might play a role in extending the interface between the globular domain of NEDD8 and CSN5, thereby, enhancing their basal interaction. In return, the strengthened binding between CSN5 and the N8∼CRL1 conjugate might stabilize the inhibitor and prolongs its retention at the CSN5 active site. Such a mechanism provides an alternative explanation for its substrate-dependent potency of the orthosteric inhibitor.

To test this idea, we first assessed the potency of CSN5i-3 in inhibiting the intrinsic catalytic activity of the CSN5-CSN6 heterodimer towards NEDD8-rhodamine. Remarkably, the CSN inhibitor blocked the reaction with an IC_50_ of 145 nM (**Extended Data Fig. 3g**). Such a high potency is in stark contrast to its μM affinity of binding free CSN5-CSN6 and is close to its potency in inhibiting N8∼CRL1 deneddylation by the intact CSN (**Fig. 3e, Extended Data Fig. 3b, c**). The unique discrepancy between the high potency and the moderate affinity of the compound towards CSN, therefore, can be recapitulated with the much-simplified enzyme-substrate system and is largely independent of CRL1. In support of this conclusion, CSN5i-3 can also inhibit the hydrolysis of NEDD8-rhodamine by CSN with a 95 nM potency ((**Extended Data Fig. 3h**).

We next asked whether the inhibitor can indeed promote the enzyme-substrate interaction as a molecular glue. Because CSN5i-3 physically occludes the CSN5 active site and does not directly contact the iso-peptide bond formed between NEDD8 and CUL1 Lys-720, we replaced NEDD8∼rhodamine with native NEDD8 and measured its affinity toward the CSN5-CSN6 heterodimer in the presence and absence of CSN5i-3. Interestingly, while the interaction between NEDD8 and CSN5-CSN6 was barely detectable by BLI, they acquired a high affinity of ∼930 nM when CSN5i-3 was present at a saturating concentration (**Fig. 5a**). Importantly, the same observation was made with the intact CSN, which can capture free NEDD8 with an ∼130 nM affinity in an inhibitor-dependent manner (**Fig. 5b**). Together, these data allowed us to conclude that the orthosteric inhibitor can indeed act as a molecular glue to seize NEDD8 conjugated to CUL1 while excluding the iso-peptide conjugate from the catalytic cleft.

**Fig. 5.**
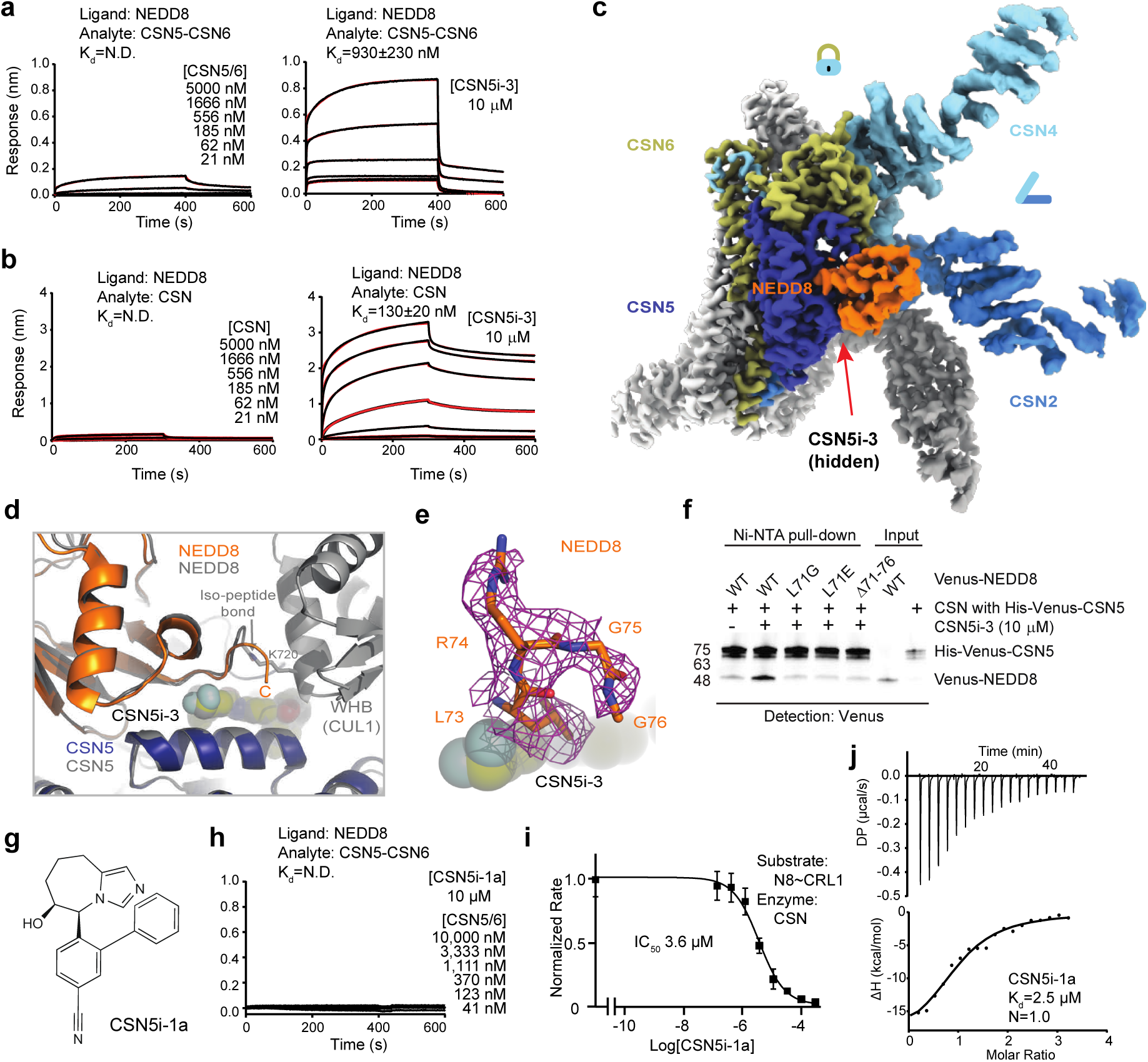
CSN5i-3 acts as a molecular glue. **a, b** NEDD8 binding to CSN5-CSN6 or CSN in the absence and presence of CSN5i-3 determined by BLI. **c.** Cryo-EM map of the CSN5i-3-CSN-NEDD8 complex. **d.** A comparison of the NEDD8 binding modes to CSN5 (left) in the CSN5i-3-CSN-N8∼CRL1 and CSN5i-3-CSN-N8 complexes with CSN5 superimposed. **e.** Well-resolved density of the NEDD8 C-terminal tail stabilized by CSN5i-3 in the CSN5i-3-CSN-NEDD8 complex. **f.** The impact of NEDD8 C-terminal tail truncation and mutations on CSN5i-3-enhanced NEDD8-CSN interactions. The protein samples were not boiled before SDS-PAGE analysis. **g.** The chemical structure of CSN5i-1a. **h.** CSN5i-1a shows no activity in promoting NEDD8 binding to CSN. **i.** The potency of CSN5i-1a in inhibiting the iso-peptidase activity of CSN towards N8∼CRL1 in vitro. **j.** Binding affinity of CSN5i-1a to the CSN5-CSN6 heterodimer determined by ITC.

To further validate the conclusion, we determined the cryo-EM structure of CSN5i-3-bound CSN in complex with NEDD8 at a resolution of 3.3 Å (**Extended Data Fig. 4a**, **Extended Data Table 1**). As expected, without CRL1 loaded, CSN in the complex adopts the same overall topology as revealed for its apo and CSN5i-3 bound forms (**Fig. 5c**). The presence of the orthosteric inhibitor, however, stabilizes the interaction between CSN and NEDD8, which becomes well resolved in the cryo-EM map with clear side chain densities (**Extended Data Fig. 4a**). A superposition analysis of the CSN5i3-CSN-N8 and CSN5i3-CSN-N8∼CRL1 structures unveils a nearly identical binding mode of NEDD8 on CSN5, albeit the differential orientations of the iso-peptidase subunit relative to the rest of the complex (**Fig. 5d, Extended Data Fig. 4b**). Although the C-terminal tail of NEDD8 is excluded from the CSN5 catalytic cleft, it is shaped into a specific coiled conformation by CSN5i-3 (**Fig. 5e**). Remarkably, removal or single amino acid mutations of the NEDD8 C-terminal tail effectively abrogated the enhanced NEDD8-CSN interaction fostered by the inhibitor (**Fig. 5f**).

CSN5i-1a is a precursor of CSN5i-3 and lacks the difluoromethyl pyrazole moiety that can directly interact with the NEDD8 C-terminal tail (**Fig. 5g**). Analogous to the NEDD8 C-terminal tail mutants, CSN5i-1a fails to promote the binding of NEDD8 to CSN (**Fig. 5h**). Moreover, it inhibits CSN only with a single digit μM potency, even though it docks to the CSN catalytic module with an affinity and binding pose similar to CSN5i-3 (**Fig. 5i,j, Extended Data Fig. 4c**). The distinction between the two compounds is further manifested by their differential effects on the binding affinity of N8∼CRL1 to the deneddylase complex they orthosterically inhibit. In the presence of a saturating amount of compound, CSN interacts with its substrate with a K_D_ of 1.8 μM and 26 nM, when being blocked by CSN5i-1a and CSN5i-3, respectively (**Extended Data Fig. 4d, e**). Together, these results reinforce the notion that CSN5i-3 strengthens the otherwise weak interaction between NEDD8 and CSN5 as a molecular glue and attains its high potency by leveraging the cooperativity of the inter-molecular interactions at the tri-molecular junction.

### Alteration of CSN interactome by CSN5i-3

As an uncompetitive inhibitor, CSN5i-3 shows a nearly irreversible effect in the cell, analogous to a covalent inhibitor ^24^. Such a property suggests that the molecular glue effect of the compound might alter the dynamic of CSN-CRL interactions in the cell, which in turn affects its dissociation from the target. To assess the impact of CSN5i-3 on the interactome landscape of CSN, we employed *in vivo* cross-linking-assisted affinity purification mass spectrometry to map the endogenous CSN-centric protein-protein interactions in the native cellular environment^31^. Two HEK293 stable cell lines that individually express HBTH-tagged CSN2 and CSN6 were utilized to isolate endogenous CSN complex and its interacting partners^32^. To capture both stable and dynamic CSN-interacting proteins, we performed *in vivo* chemical cross-linking prior to cell lysis. The purified protein complexes were digested and analyzed by liquid chromatography tandem mass spectrometry (LC MS/MS) with both data-dependent (DDA) and data-independent (DIA) acquisitions. These analyses allowed us to identify not only all cullin proteins, RBX1/2, and all cullin adaptors (SKP1, ELONGIN-B/ELONGIN-C, and DDB1), but also 159 CRL substrate receptors that are associated with CSN in the cell (**Fig. 6a, Extended Data Table 3a, b**).

**Fig. 6.**
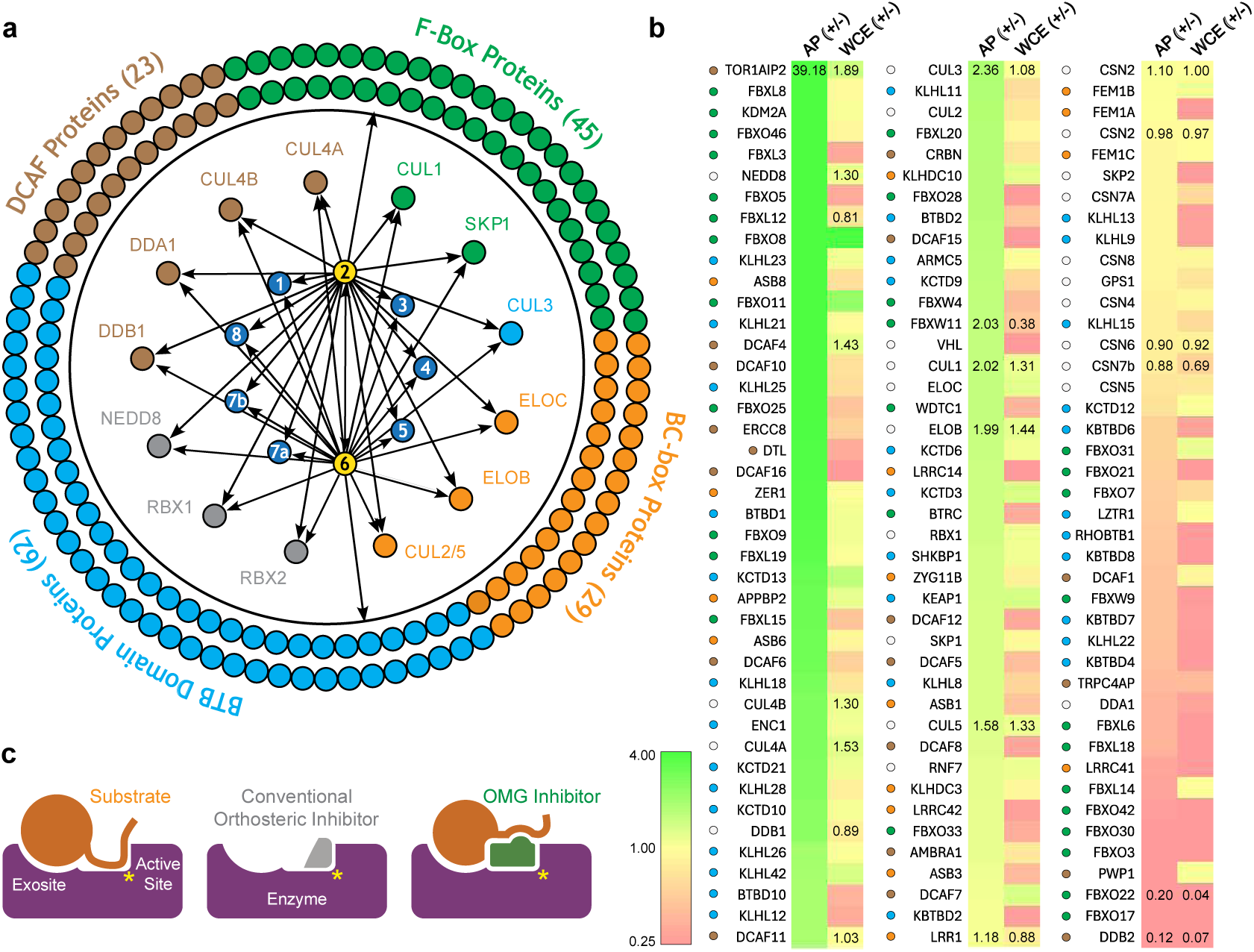
CSN5i-3-induced changes in the CSN interactome. **a.** CSN interaction network assessed by MS analyses of affinity purified CSN2 and CSN6 complexes. The central numbered circles represent the eight subunits of CSN. The CRL SR profiles for CSN2 and CSN6 affinity purification are indicated by a dash line. **b.** CSN5i-3-induced abundance changes of selected proteins in affinity-purified (AP) CSN complexes and whole-cell extracts (WCE) determined by quantitative MS analyses. “+/−”: treated (CSN5i-3)/untreated. **c.** Schematic drawing of the mechanism of action of an orthosteric moleucular glue inhibitor vs. conventional orthosteric inhibitor.

We next performed DDA- and DIA-based label-free quantitative analyses to compare the levels of CSN subunits and their interacting proteins in the purified complexes from CSN5i-3-treated and untreated cells (**Extended Data Table 3a, b**). In addition, we carried out parallel reaction monitoring (PRM)-based targeted quantitation to examine protein expression levels of CSN subunits, CRLs and 101 SRs in cells before and after CSN5i-3 treatment (**Extended Data Table 3c**). As expected, the amount of CSN subunits remained largely constant, indicating that the integrity of the iso-peptidase complex was not perturbed by the compound (**Fig. 6b, Extended Data Table 3d**). Although CSN5i-3 treatment induced minimal changes in the overall abundance of all cullins and NEDD8 in cells, it increased their association with CSN (**Fig. 6b, Extended Data Table 3d**). This could be attributable to the accumulation of N8∼CRLs in the cell and possibly the inhibitor-induced stabilization of the enzyme-substrate assembly. Strikingly, the amount of different CRL substrate receptors copurified with CSN exhibited a broad range of CSN5i-3-induced changes, varying from near-complete depletion to pronounced enrichment. At one end of the spectrum, a fraction of CRL substrate receptors, such as DDB2 and FBXO22, showed diminished interactions with CSN upon CSN5i-3 treatment (**Fig. 6b**). Their total protein level, however, dropped prominently in the cell. These observations were confirmed by immunoblotting analysis (**Extended Data Fig. 4f-h**). The overall destabilization of these CRL substrate receptors, therefore, corroborates with the proposed role of CSN in protecting them from auto-ubiquitination and subsequent degradation. At the other side of the spectrum, it is surprising that the majority of CRL substrate receptors such as DCAF4 and FBXL12 showed elevated association with CSN upon inhibitor treatment even though their expression levels remained largely unchanged or was decreased (**Fig. 6b, Extended Data Fig. 4f**). This phenomenon, while contradictory to the hypothesized function of CSN, can be explained by the molecular glue activity of CSN5i-3 in locking the enzyme-substrate complexes. Previous structural analyses have revealed that CSN subunits outside the catalytic hemisphere can make direct contacts with select CRL substrate receptors^4^. It is plausible that these interfaces can synergize with CSN5i-3 to differentially trap N8∼CRLs with substrate receptors on CSN.

## DISCUSSION

Orthosteric inhibitors are small molecules that compete with a substrate for binding to the active site of an enzyme, whereas molecular glues are compounds that enhance protein-protein interactions by extending and stabilizing their interfaces. At face value, these two classes of agents appear to operate through opposing, if not mutually exclusive, mechanisms of action. However, when the substrate of an enzyme is a protein, a combined mechanism of action becomes possible. This is because the substrate and the enzyme can interact through a nearby exosite distinct from the active site. In this study, we discovered that CSN5i-3, a potent CSN inhibitor, functions dually as an orthosteric inhibitor of the iso-peptidase target while simultaneously stabilizing the enzyme-substrate complex as a molecular glue. Such an unexpected mechanism explains how the compound inhibits N8∼CRL deneddylation by CSN uncompetitively and gains its high potency in a substrate-dependent manner. Based on this finding, we propose a new concept of orthosteric molecular glue (OMG) inhibitors (**Fig. 6c**).

As an OMG inhibitor, CSN5i-3 possesses two unique properties that distinguish it from conventional orthosteric enzyme inhibitors. First, CSN5i-3 binds to the active site of its target with a moderate μM affinity but can inhibit the iso-peptidase complex with a low nM high potency. The apparent discrepancy of the two parameters is attributed to the presence of the substrate when the enzymatic activity is determined. The high potency of the OMG inhibitor stems from the cooperativity of inter-molecular interactions centered around the molecular glue molecule at the enzyme-substrate interface. While a molecular glue compound enhances the basal affinity between two weakly interacting proteins, the resulting protein complex formation can in turn stabilize the small molecule trapped at the protein interface and reduce its off rate. Such a special attribute of OMG inhibitors, therefore, points to an immediate implication in drug discovery. For instance, tailoring a canonical high affinity orthosteric enzyme inhibitor to an OMG inhibitor with a substrate-dependent high potency could potentially overcome its off-target toxicity.

Second, while blocking N8∼CRL1 from accessing the active site of CSN, CSN5i-3 makes direct contacts with the substrate proteins to stabilize the enzyme-substrate assembly. This unexpected property affords a unique opportunity for an OMG inhibitor to achieve selectivity towards different substrates of the enzyme it targets, as long as there are variations of substrate sequence at the substrate-OMG inhibitor interfaces. In the case of CSN5i-3, its molecular glue activity can be affected by a single amino acid in NEDD8 and possibly by the CSN-CRL substrate receptor interface. Even if the interfaces between different substrates and an OMG inhibitor is invariant, differential PPI strength at the exosite could conceivably confer unequal potency of the compound in inhibiting the same enzyme towards different substrates. Such a special feature can be particularly valuable for enzymes, whose disease-relevant versus toxicity-related substrates might be expressed in different tissues.

The mechanism of action of CSN5i-3 is somewhat reminiscent of selected inhibitors of enzymes, such as CDK12, MEK1/2, KIF18A, and BRD4, that bind to either the active site or an allosteric pocket of their targets and promote their interactions with other proteins^11,12,33–39^. Nevertheless, CSN5i-3 is distinct from these enzyme inhibitors and other known molecular glues by functioning as an orthosteric enzyme inhibitor with substrate-dependent potency. Its unique features echo the methylthioadenosine (MTA)-dependent PRMT5 inhibitors and showcase how cooperativity of molecular interactions can drive context-dependent enzyme inhibition^40^. While the discovery of CSN5i-3 as an OMG inhibitor was fortuitous, we envision that CSN5i-3-like OMG inhibitors can be developed by design for many clinically relevant enzyme targets that recognize substrates proteins via an exosite adjacent to the catalytic pocket. These include intracellular and extracellular proteases, and many enzymes catalyzing the forward and reverse reactions underlying protein post-translational modifications. Monitoring the impact of derivative compounds of a canonical orthosteric inhibitor on the enzyme-substrate interaction while improving their potency can guide the development of an OMG inhibitor.

## Data and code availability

The coordinates and cryo-EM maps were deposited in the Protein Data Bank (PDB) and the Electron Microscopy Data Bank (EMDB) with the following accession numbers: CSN: 9E5Z, EMD-47532; CSN5i-3-CSN: 9E81, EMD-47698; CSN5i-1a-CSN: 9PH4, EMD-71639; CSN5i-3-CSN-N8∼CRL1: 9EFV, EMD-47981, EMD-47729, EMD-47767; -CSN5i-3-CSN-NEDD8: 9E77, EMD-47660; CSN^DM^-N8∼CRL1: 9EFM, EMD-47976, EMD-47500, EMD-47502; CSN^DM^-N8∼CRL2: 9EFQ, EMD-47977, EMD-47701, EMD-47702; CSN^DM^-N8∼CRL3: 9EGL, EMD-47990, EMD-47776, EMD-47985; CSN^DM^-N8∼CRL4A: 9EG8, EMD-47986, EMD-47543, EMD-47699, and CSN^DM^-N8∼CRL5: 9EG1, EMD-47983, EMD-47663, EMD-47665.

## Acknowledgments

The authors would like to thank R. Yan, X. Zhao, J. Jung, and Z. Yu at the Cryo-EM Facility on the Janelia Research Campus of the Howard Hughes Medical Institute, and J.D. Quispe and S. Dickinson at the Arnold and Mabel Beckman Cryo-EM Center at the University of Washington for their assistance in electron microscopy data acquisition, and Paul Morenkov at University of California Irvine for his assistance on formatting TMT XL-MS data. N.Z. is Howard Hughes Medical Institute Investigators. This work is also supported by NIH grants R35GM145249 and R01CA290875 to L.H., R01AI179768 and R01AI152358 to E.F., and ISF grant 737/23 to D.S.

## Author contributions

H.S., H.M., L.H., and N.Z. conceived the project. H.M. and H.S. constructed and purified all proteins samples in this study with the help from S.C. H.S. performed all cryo-EM grid preparation, specimen screening, data collection and processing tasks. T.R.H. and H.S. performed the in vitro enzymatic and binding assays. X.W. prepared samples for MS analyses and performed biochemical experiments. C.Y. performed mass spectrometry experiments and data analysis. F.J. performed in vitro cross-linking and mass spectrometry experiments. M.B., B.S., and D.S. analyzed the in vitro cross-linking mass spectrometry data. Z.Z. and E.F. synthesized CSN5i-1a. The manuscript was written by N.Z., H.S., and L.H. with inputs from all authors.

---

**Extended Data Fig. 1.**
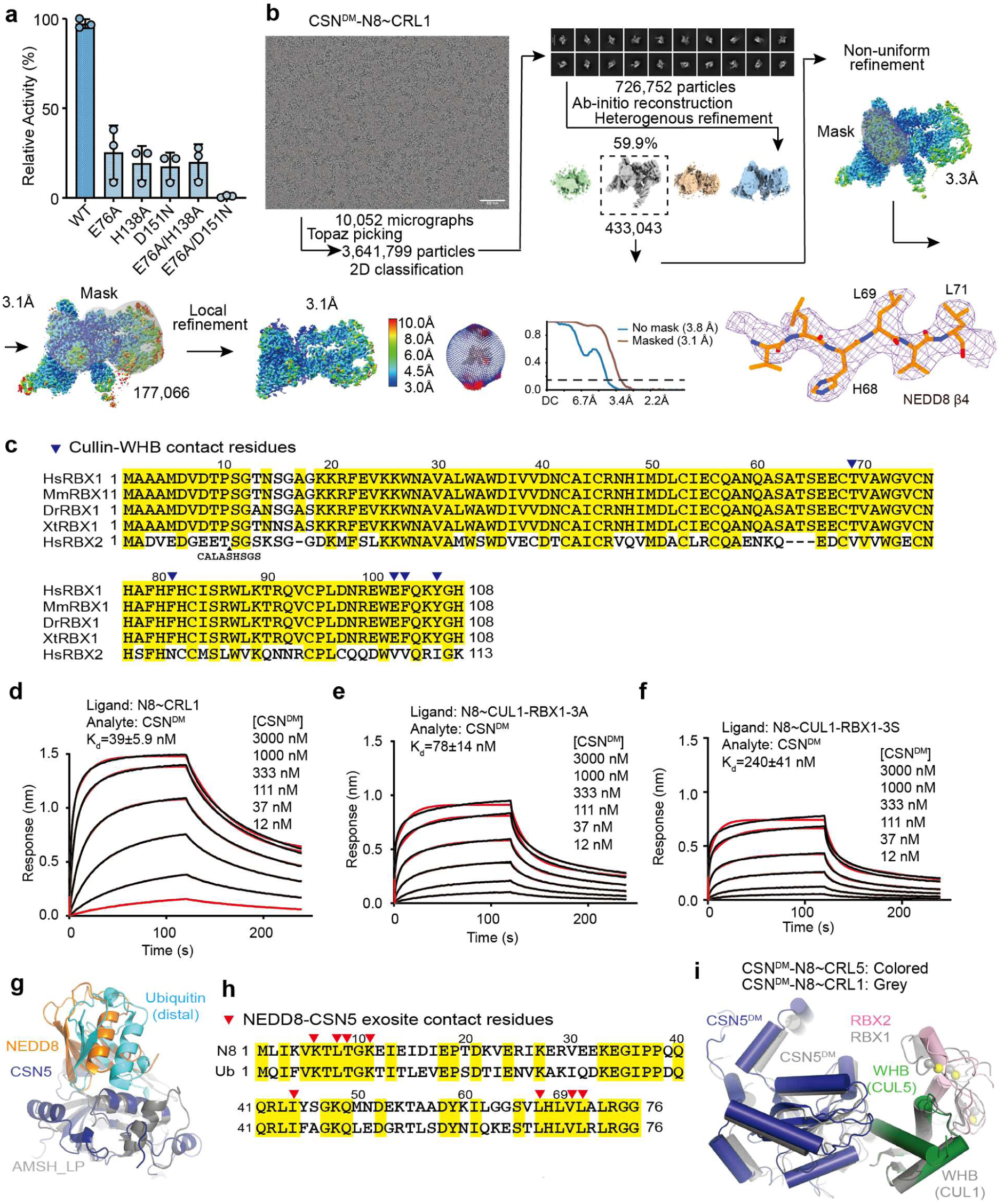
Structure determination of the pre-catalytic state of CSN. **a.** A comparison of the enzymatic activities of wild type (WT) and mutant CSN. **b.** Cryo-EM data processing for the CSN^DM^-N8∼CRL1 complex. **c.** Sequence alignment of RBX1 orthologues and human RBX2. Hs: Homo sapiens, Mm: Mus musculus, Dr: Danio rerio, Xt: Xenopus tropical. Cullin-WHB-contacting residues are indicated by blue triangles. **d-f.** Affinity measurement of CSN^DM^ to N8∼CRL1 containing wild type and mutant RBX1 by BLI. RBX1-3A and RBX1-3S feature the three aromatic residues at the RBX1-WHB interface mutated to alanine and serine, respectively. **g.** A comparison of the binding mode of NEDD8 to CSN5 in the CSN5i-3-CSN-N8∼CRL1 complex to that of the distal ubiquitin to AMSH-LP in the AMSH-LP K63-linked ubiquitin dimer complex (PDB:2ZNV). **h.** Sequence alignment of human NEDD8 and ubiquitin. CSN5-contacting residues in NEDD8 at the NEDD8-CSN5 exosite are indicated by red triangles. Conversed residues are highlighted in yellow. **i.** A slight positional shift of CSN5 in the structure of CSN^DM^ in complexes with N8∼CRL1 versus N8∼CRL5.

**Extended Data Fig. 2.**
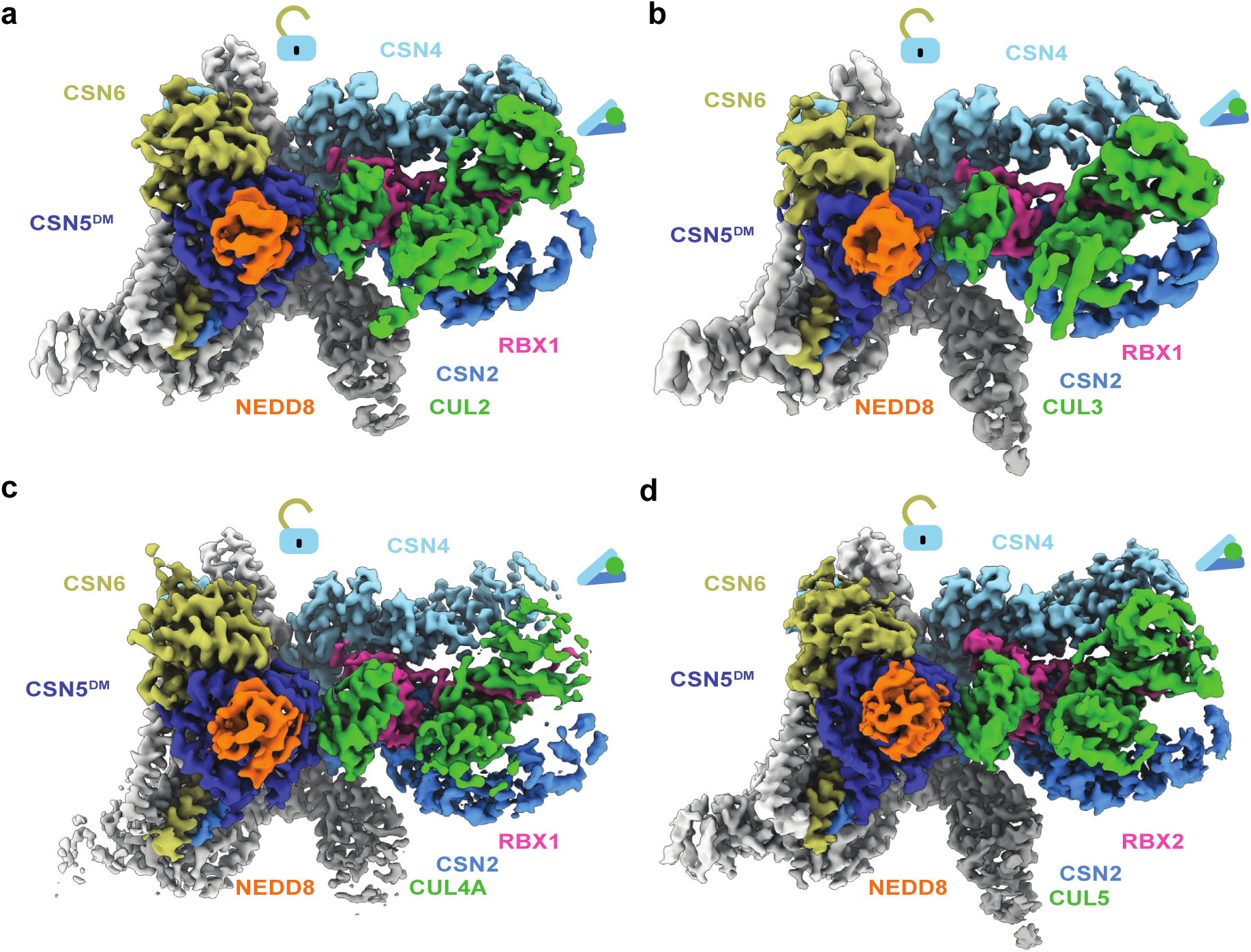
Structures of the pre-catalytic state of CSN in complexes with different N8∼CRLs. **a-d.** Cryo-EM maps of CSN bound to N8∼CRL2, N8∼CRL3, N8∼CRL4A, and N8∼CRL5.

**Extended Data Fig. 3.**
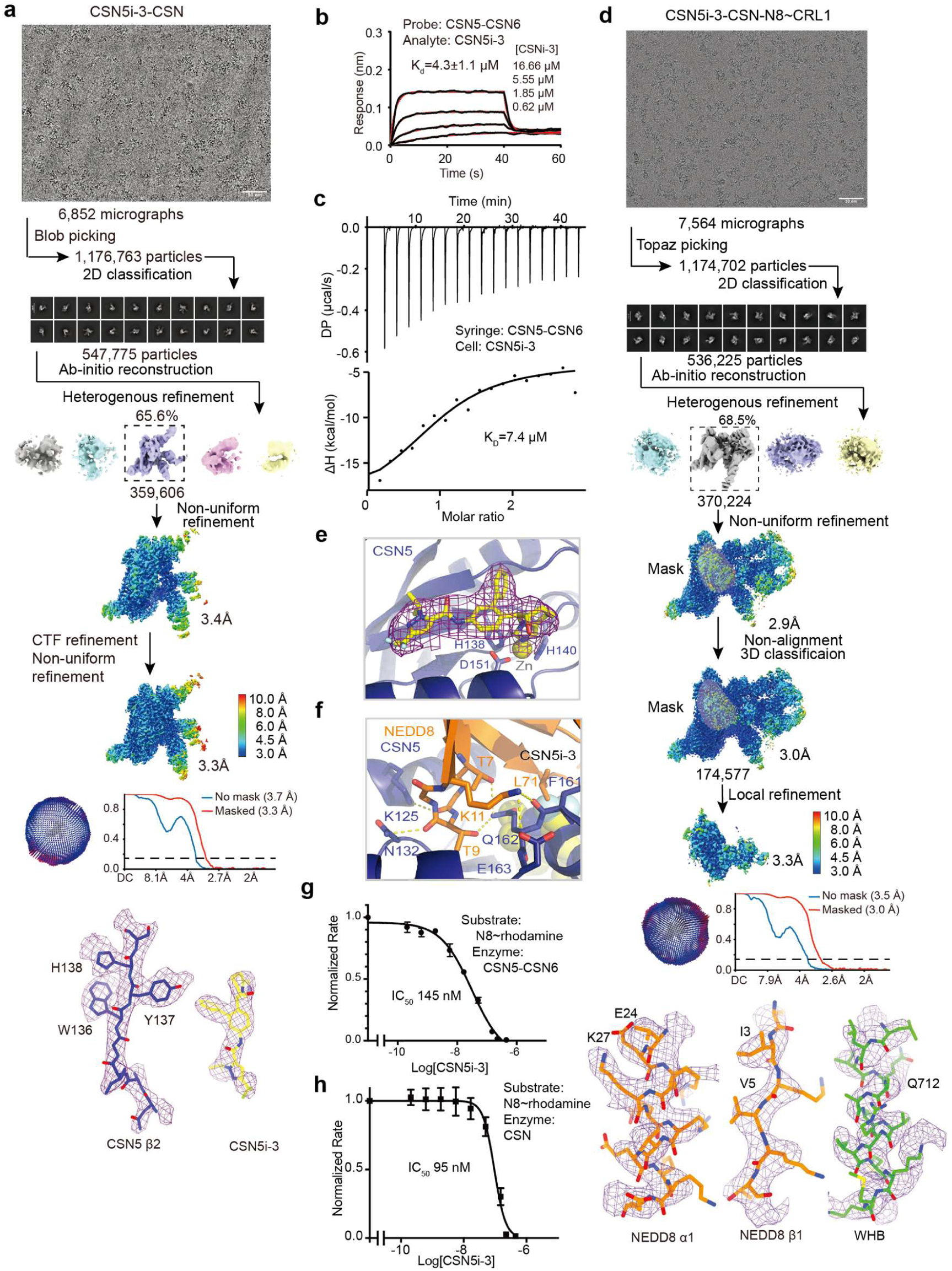
CSN5i-3 as an orthosteric inhibitor of CSN. **a.** Cryo-EM data process of the CSN5i-3-CSN complex. **b.** Binding affinity of CSN5i-3 to the CSN5-CSN6 heterodimer determined by BLI. **c.** Binding affinity of CSN5i-3 to the CSN5-CSN6 heterodimer determined by ITC. **d.** Cryo-EM data process of the CSN5i-3-CSN-N8∼CRL1 complex. **e.** A close-up view of the CSN5 catalytic site occupied by CSN5i-3 with its density revealed in the cryo-EM map of the CSN5i-3-CSN complex. **f.** A close-up view of the interface between CSN5 and the tip of the β1-β2 loop of NEDD8. Hydrogen bonds and salt bridges are shown in yellow dash lines. The side chains of interfacial residues are shown in stick. The nearby CSN5i-3 is shown in the background. **g.** The potency of CSN5i-3 in inhibiting the iso-peptide activity of the CSN5-CSN6 heterodimer towards NEDD8-Rhodamine. **h.** The potency CSN5i-3 in inhibiting the iso-peptide activity of intactCSN5 towards NEDD8-Rhodamine.

**Extended Data Fig. 4.**
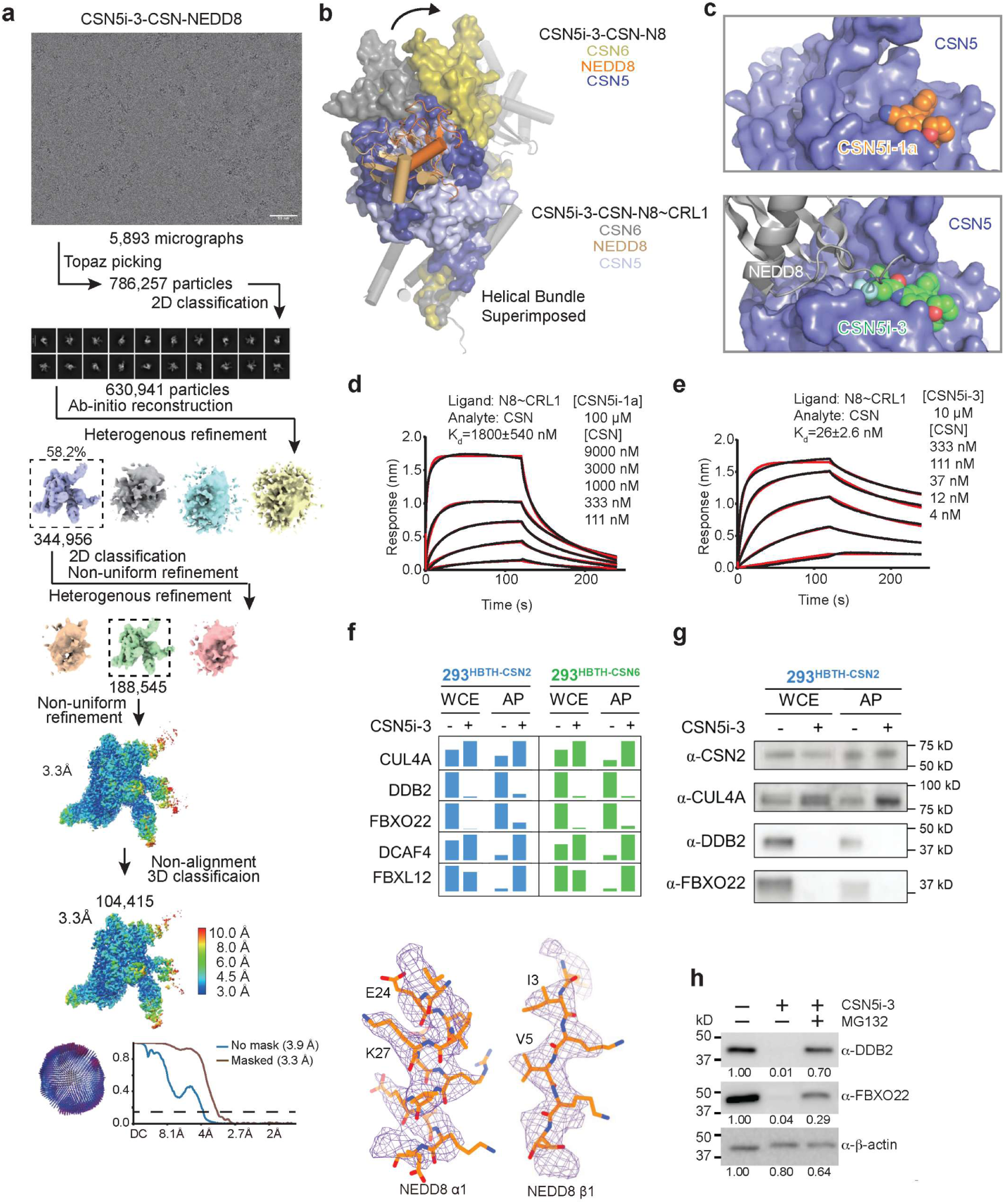
CSN5i-3 as an orthosteric molecular glue. **a.** Cryo-EM data process of the CSN5i-3-CSN-NEDD8 complex. **b.** Orientational difference of the CSN5-CSN6 dimer within CSN between the CSN5i-3-CSN-N8∼CRL1 and CSN5i-3-CSN-N8 complex structures. **c**. A comparison between the binding mode of CSN5i-1a (top panel) and CSN5i-3 (bottom panel) to CSN. Both compounds are shown in spheres. CSN5 is shown in surface representation. NEDD8 bound to CSN5i-3-occupied CSN is shown in gray cartoon. **d**. The affinity of CSN towards N8∼CRL1 in a saturating amount of CSN5i-1a measured by BLI. **e**. The affinity of CSN towards N8∼CRL1 in a saturating amount of CSN5i-3 measured by BLI. **f**. Abundances of CUL4A, DDB1, FBXO22, DCAF4 and FBXL12 in WCE and affinity-purified (AP) CSN2 and CSN6 complexes determined by quantitative MS analyses. **g.** Assessment of CUL4A, DDB1, and FBXO22 abundances in WCE and affinity-purified CSN2 complexes in the absence and presence of CSN5i-3 using immunoblotting analysis. **h**. Proteasomal degradation of DDB2 and FBXO22 induced by CSN5i-3.

**Extended Data Table 1.**
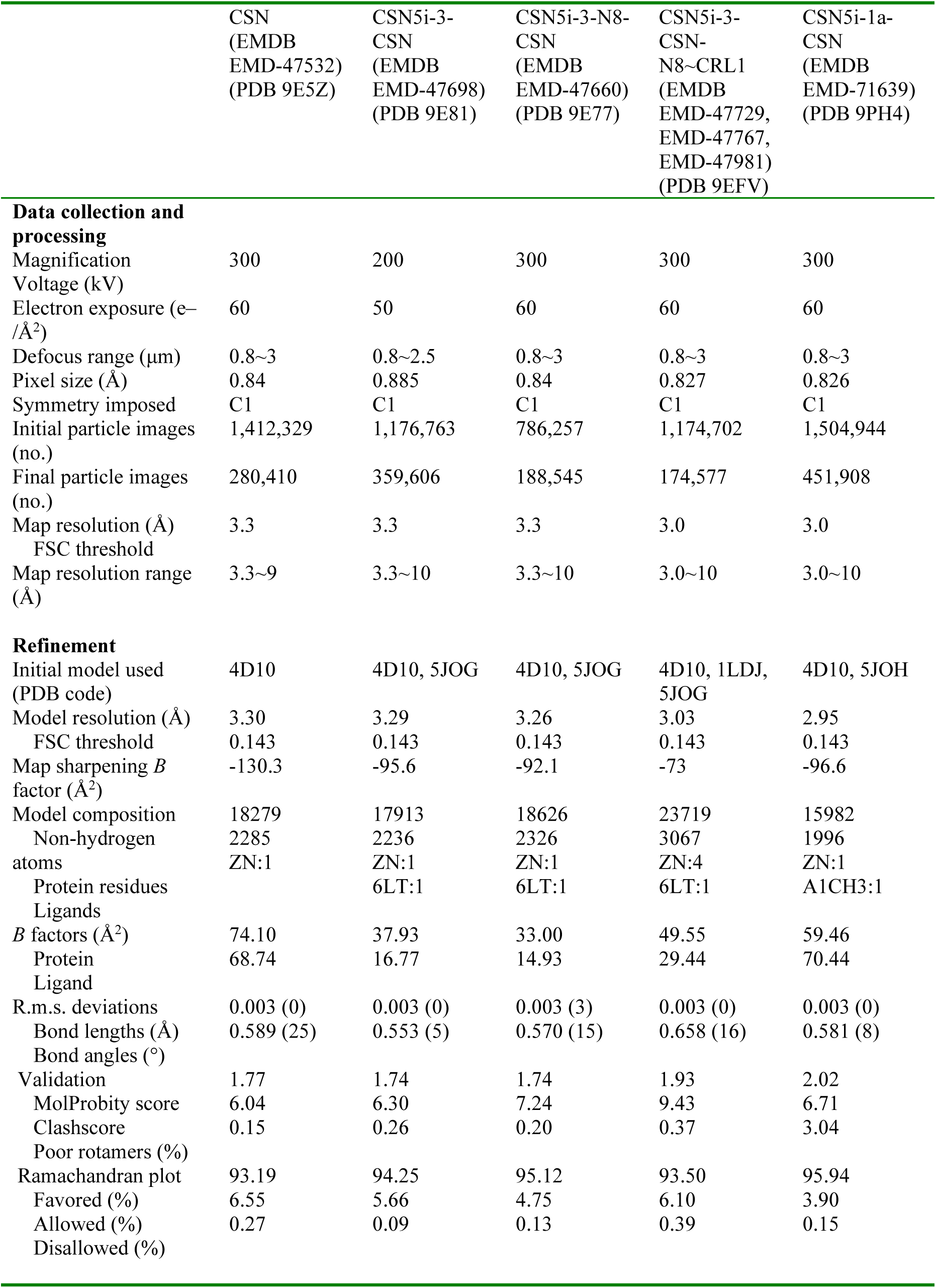
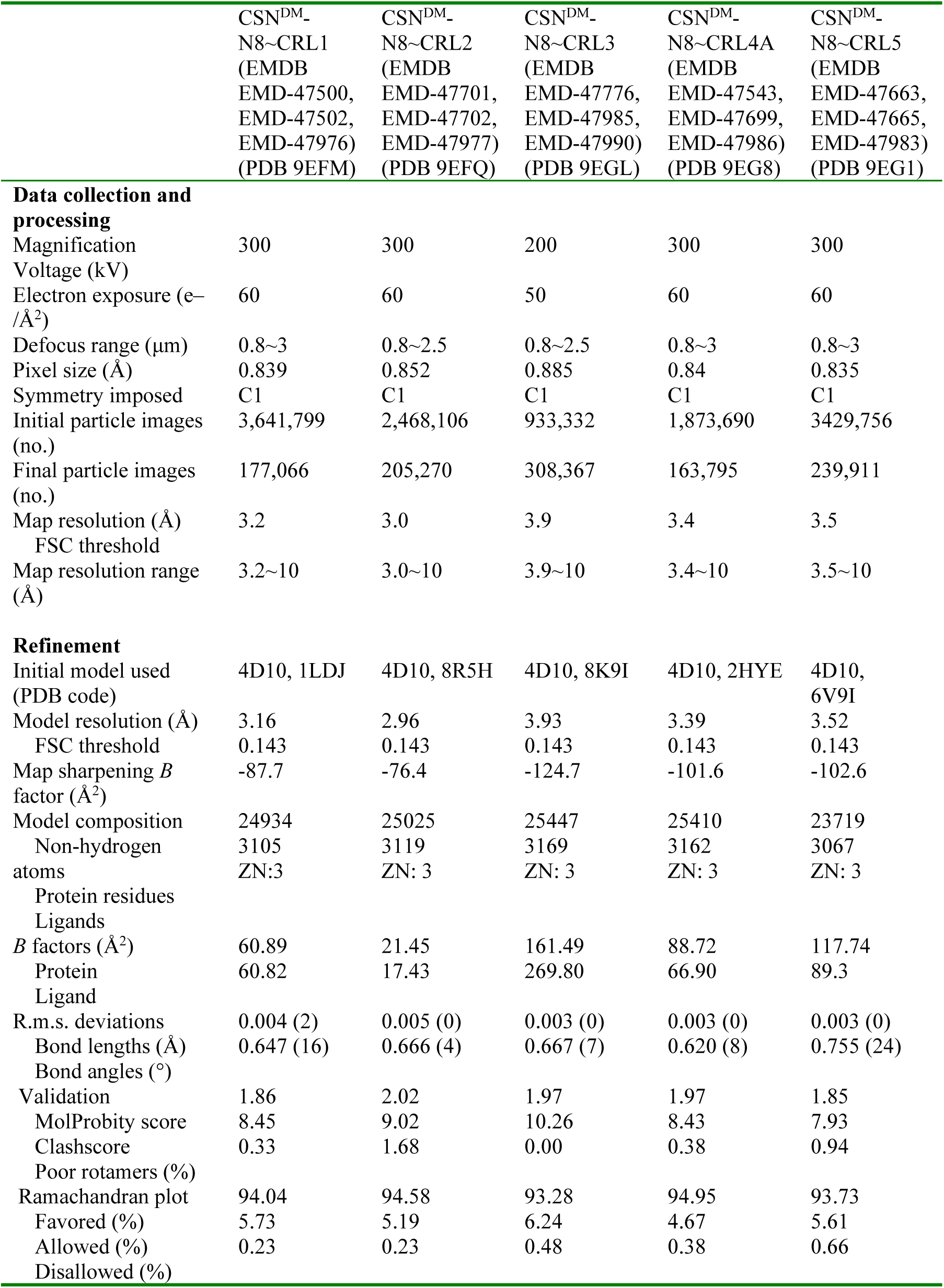
Cryo-EM image processing and model refinement statistics.

**Extended Data Table 2.**
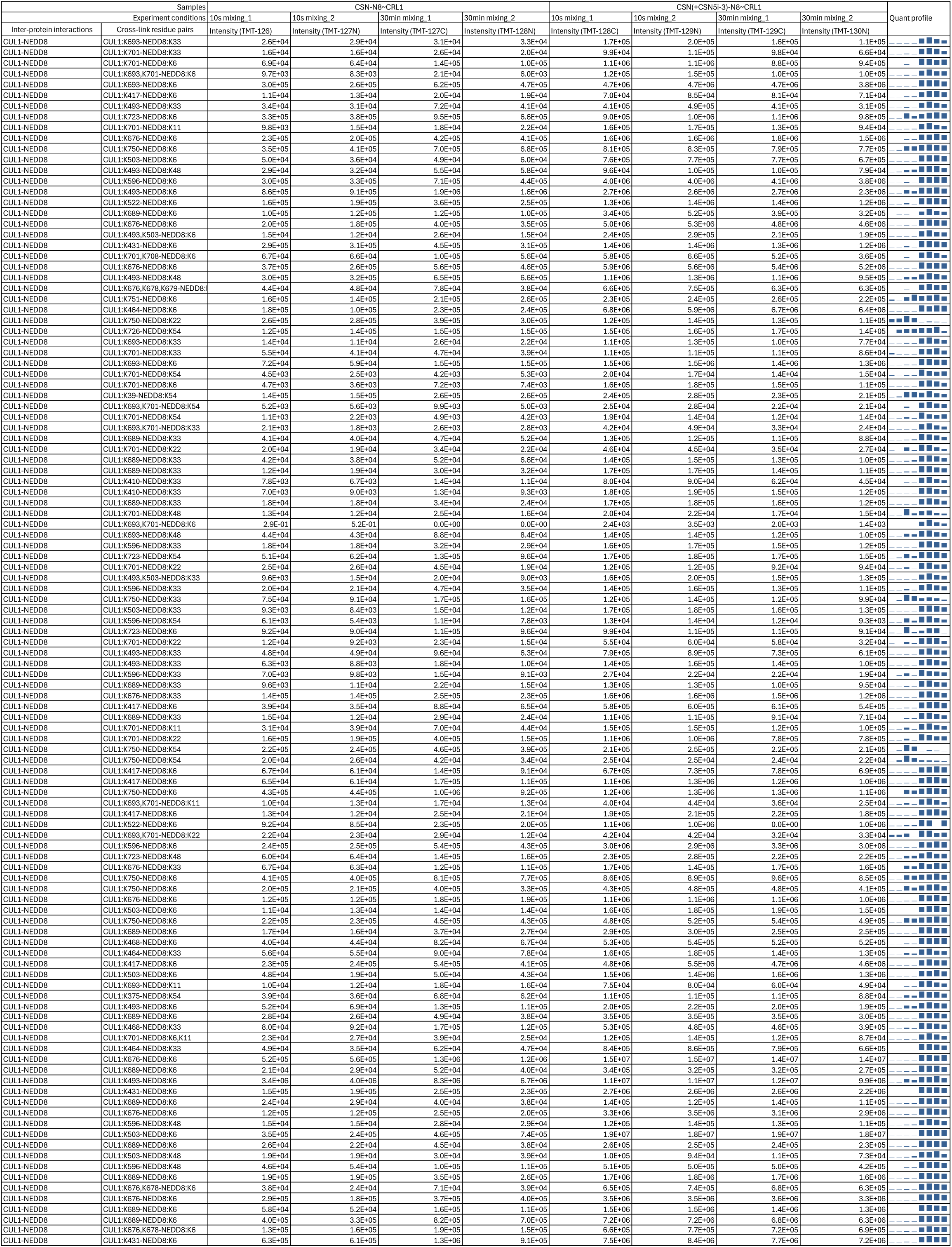
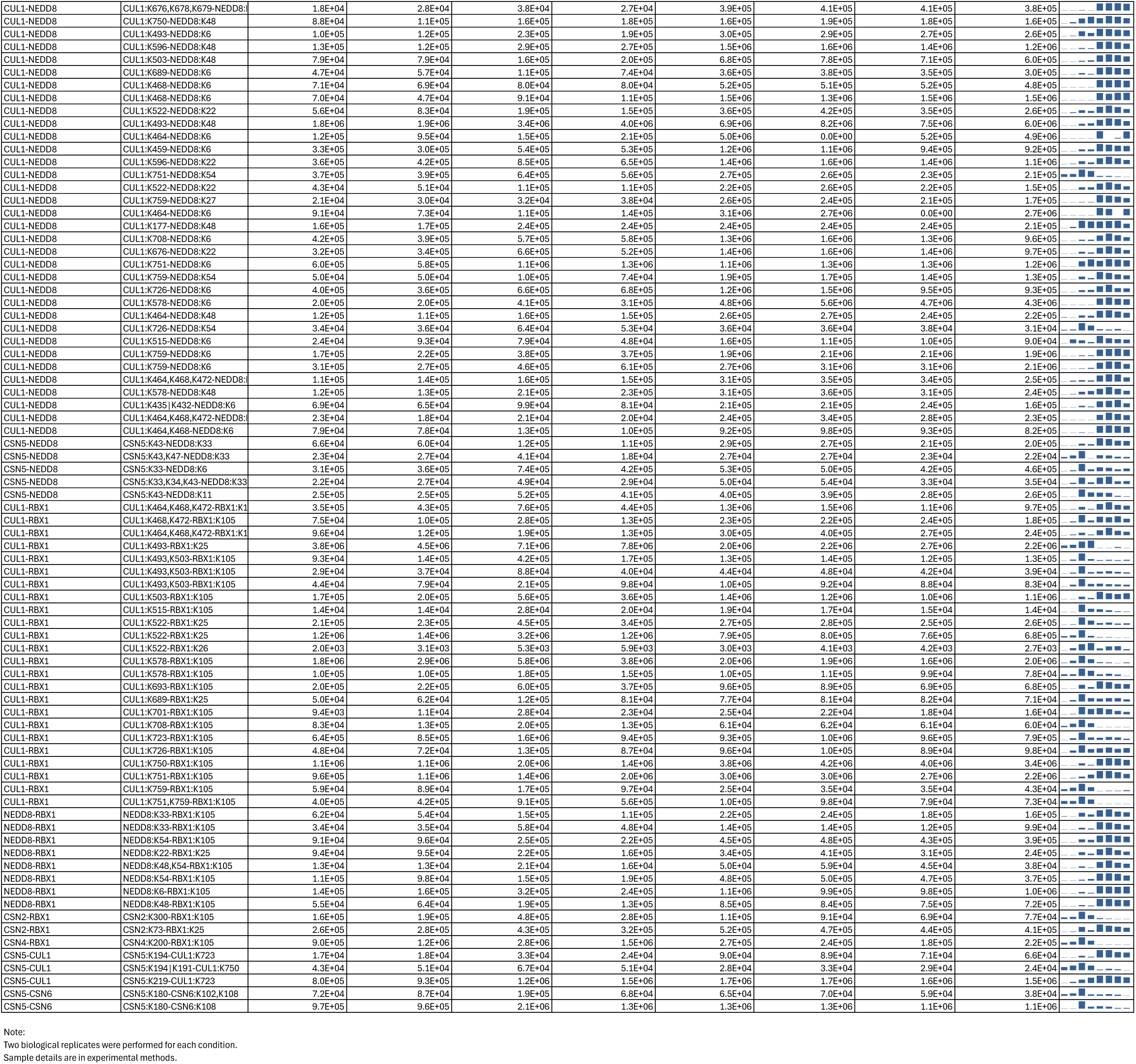
Quantitative analysis of the inter-protein cross-links of the in vitro reconstituted CSN-N8∼CRL1 complex in the presence and absence of CSN5i-3.

**Extended Data Table 3a.**
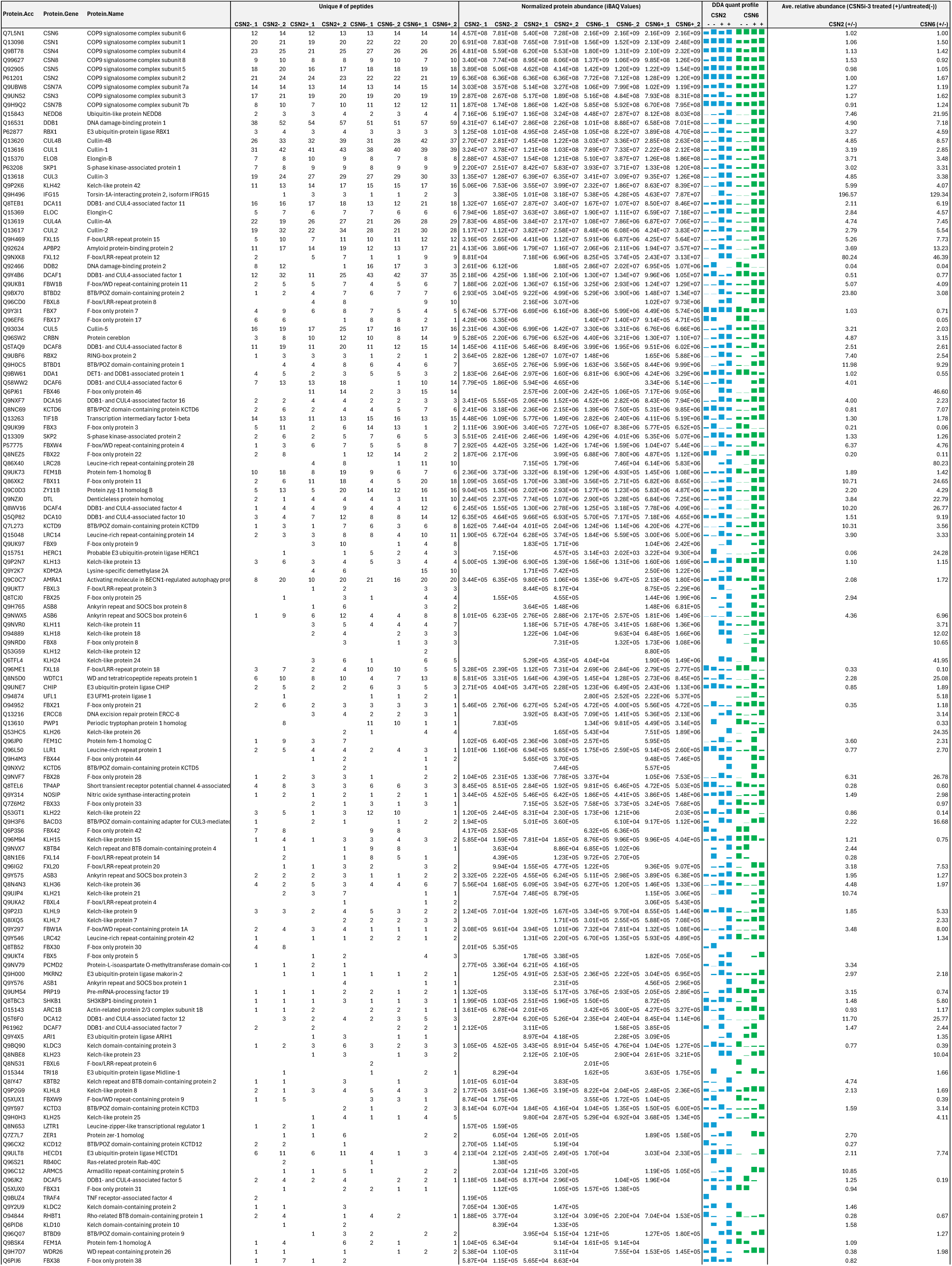
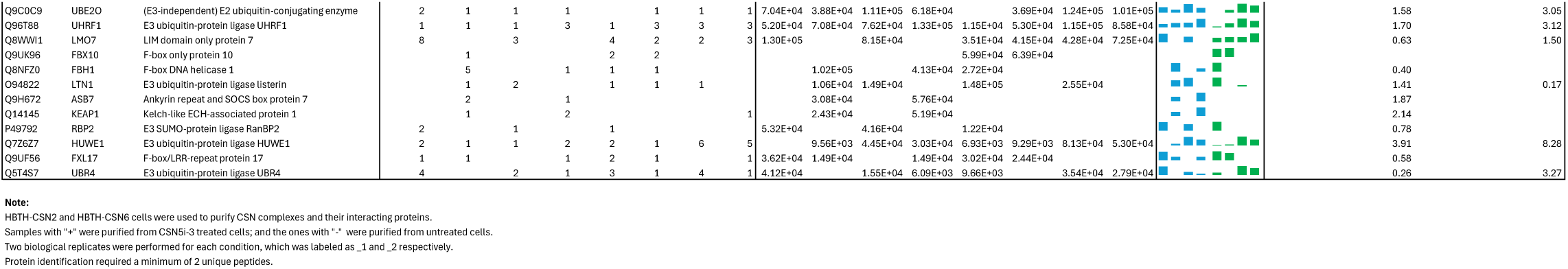
DDA-based quantitation of proteins co-purified with CSN complexes from CSN5i-3 treated and untreated cells.

**Extended Data Table 3b.**
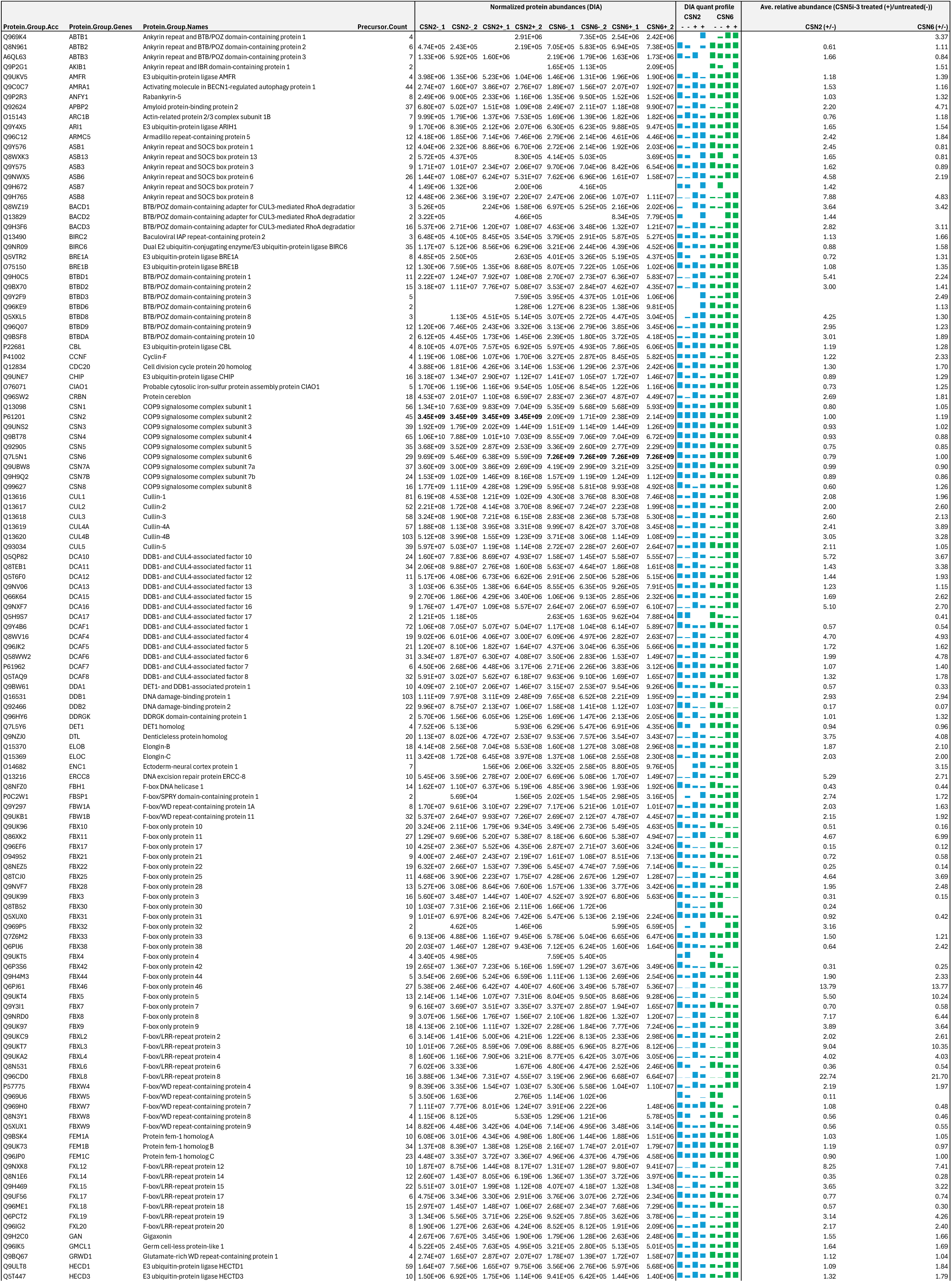
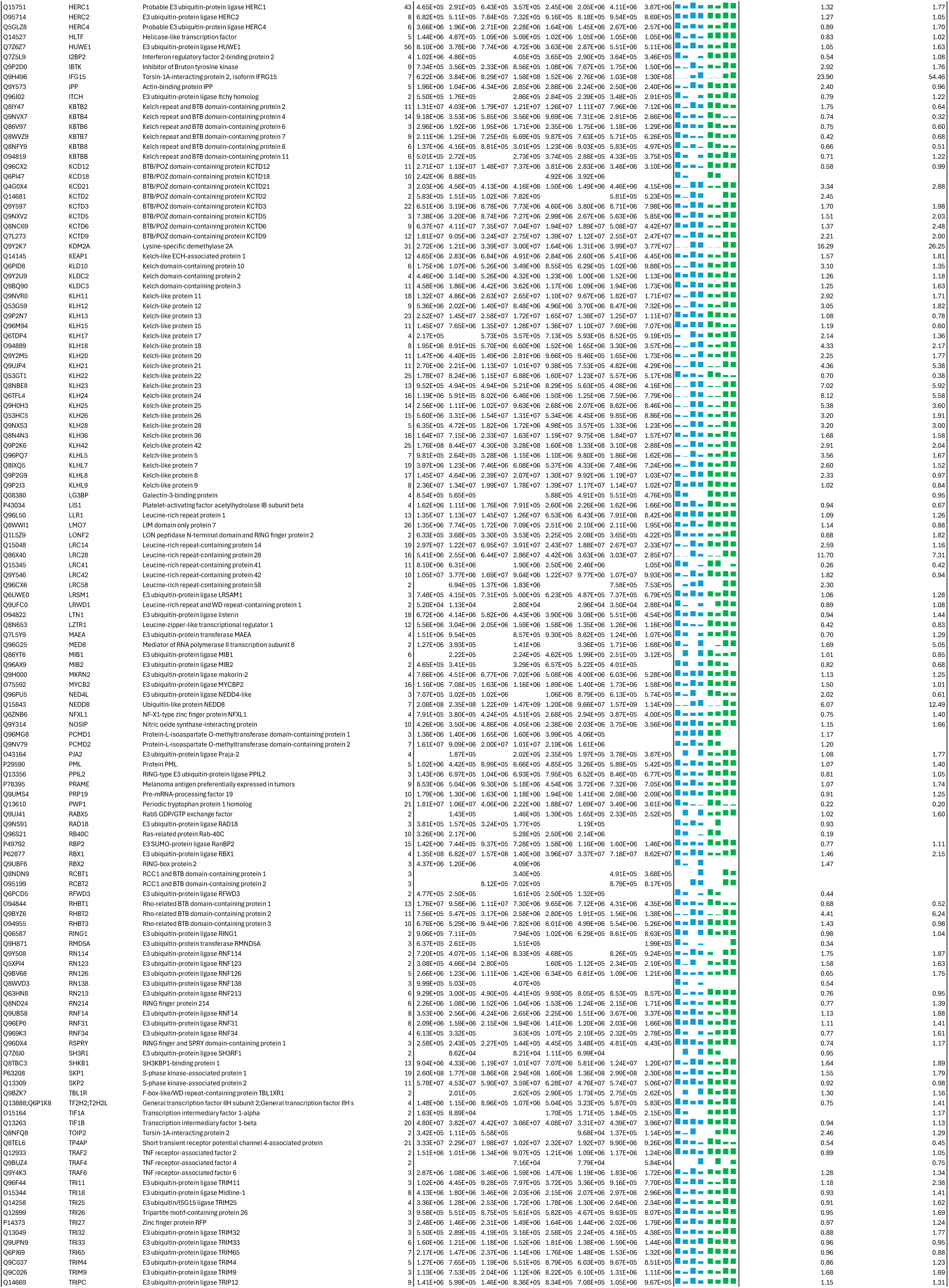
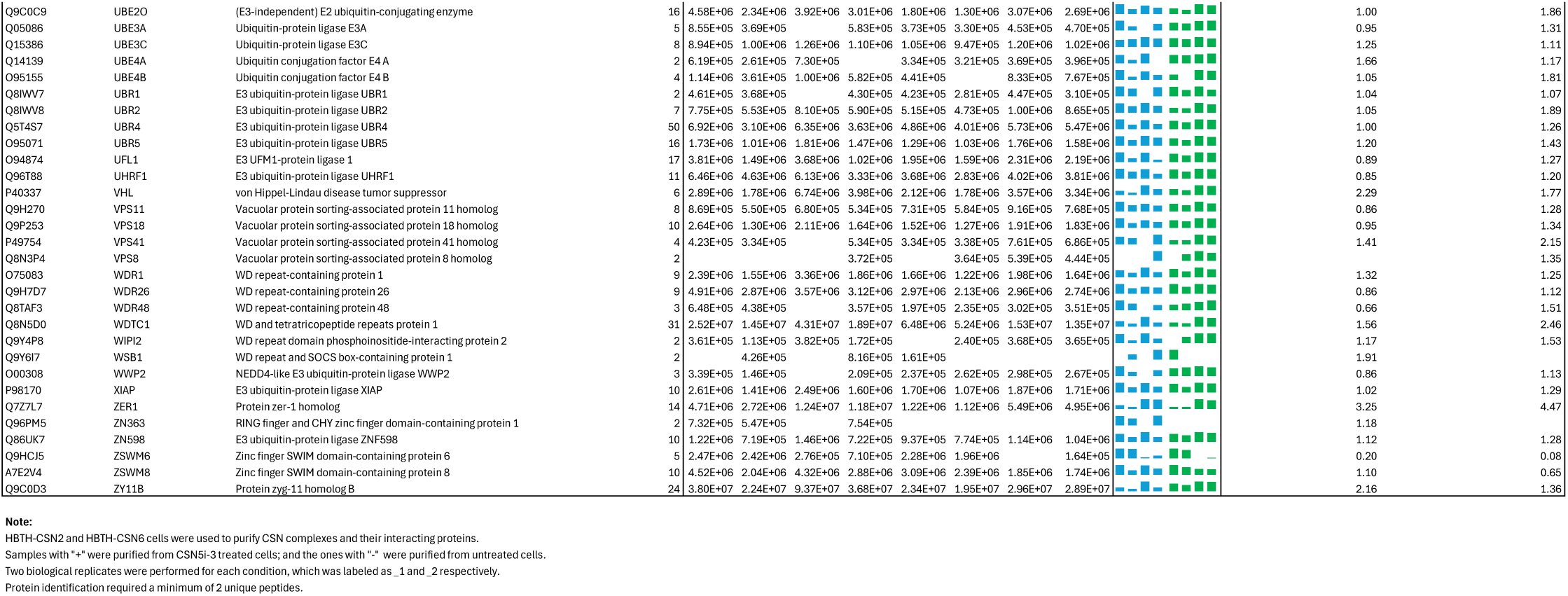
DIA-based quantitation of proteins co-purified with CSN complexes from CSN5i-3 treated and untreated cells.

**Extended Data Table 3c.**
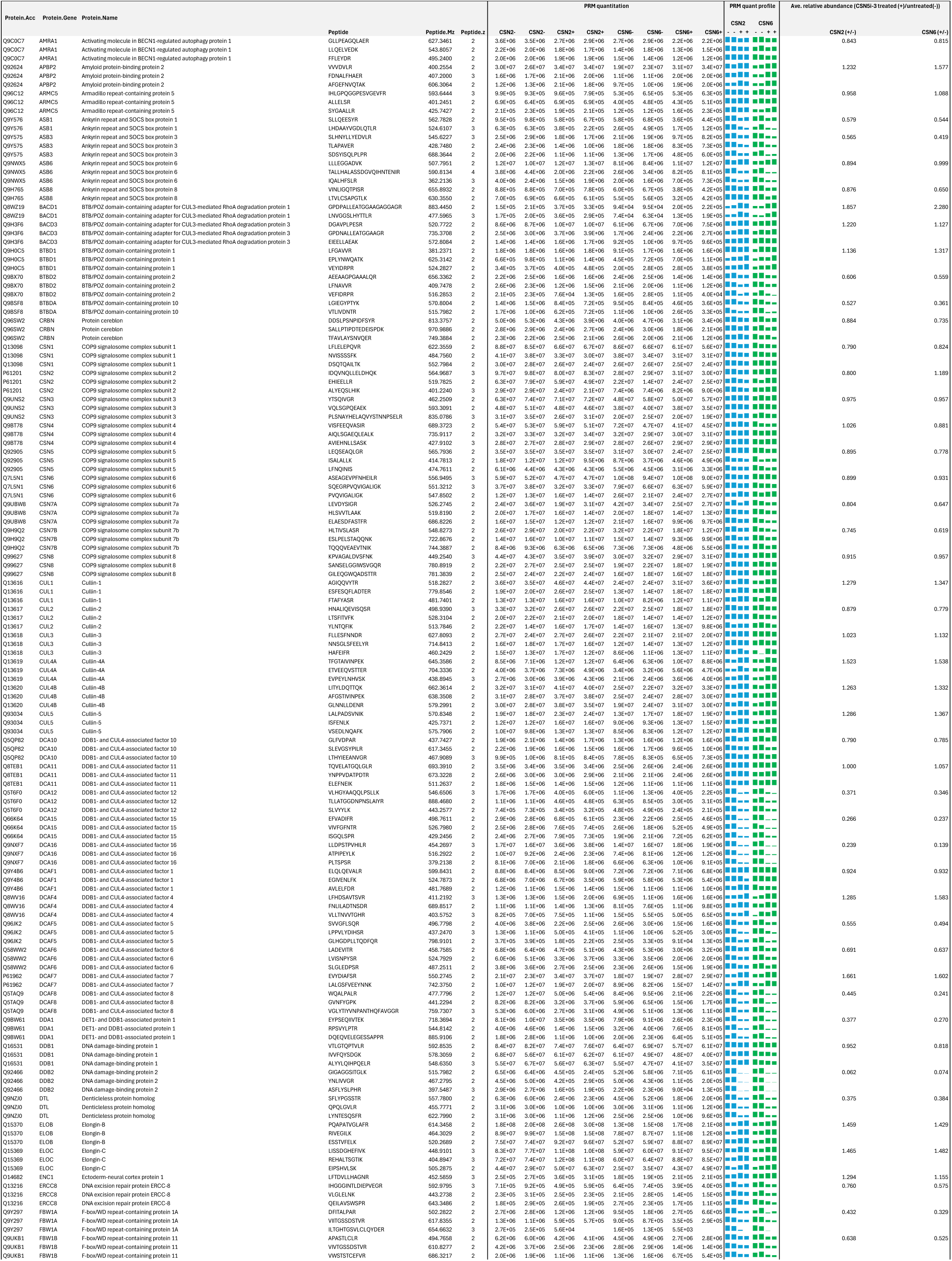
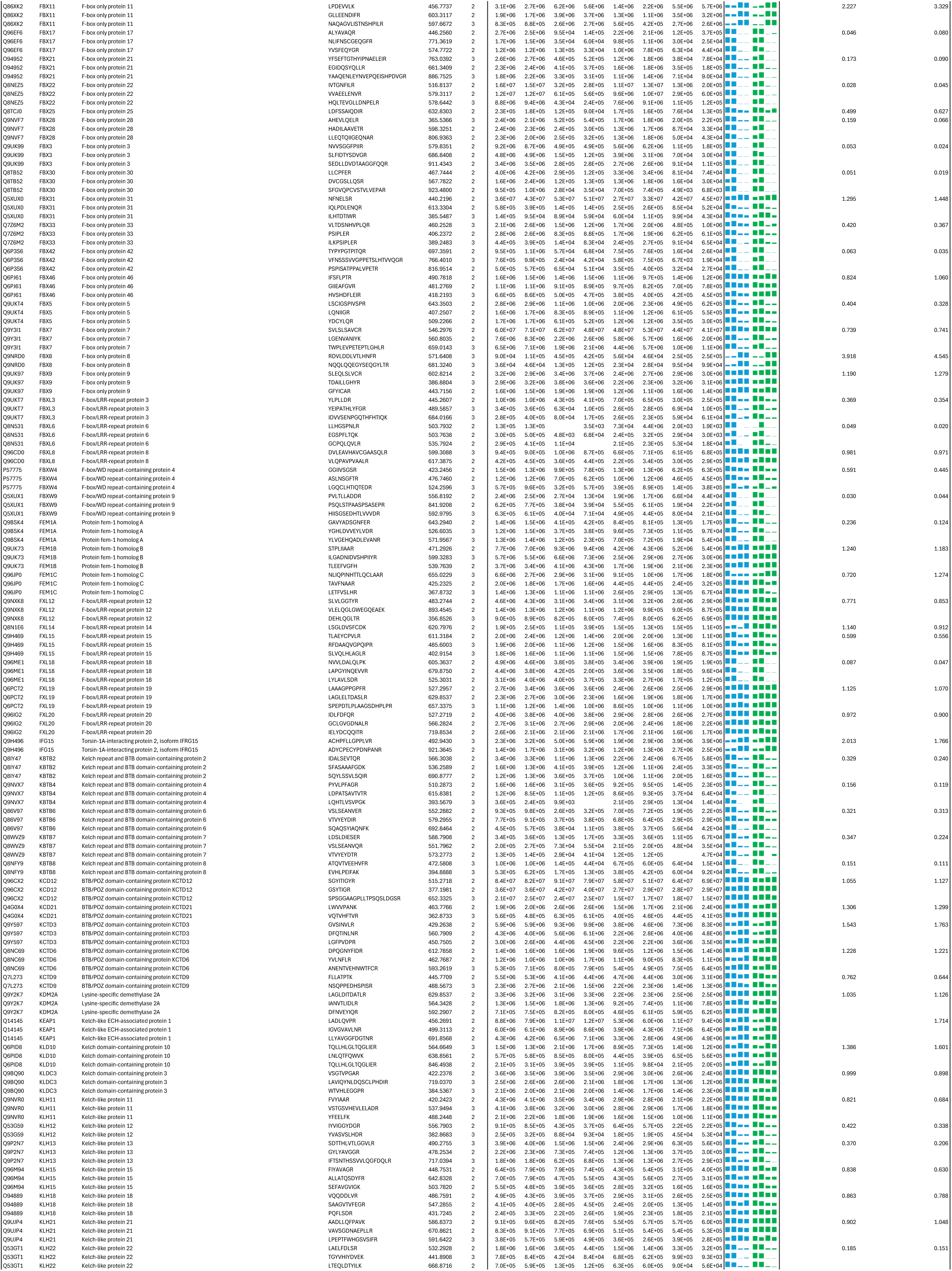
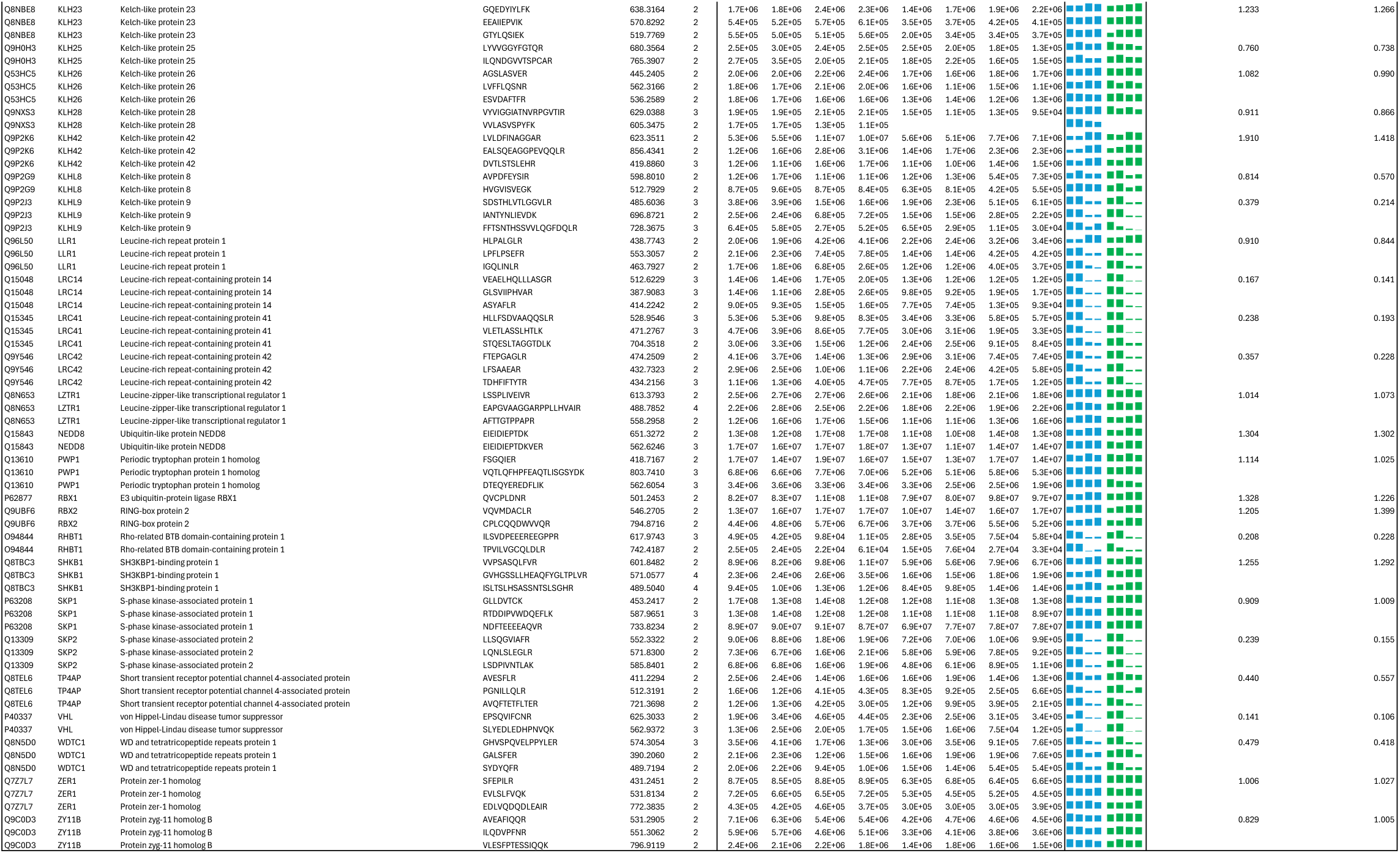
PRM-based targeted quantitation of CSN, CRLs and selected SRs in the lysates of CSN5i-3 treated and untreated cells.

**Extended Data Table 3d.**
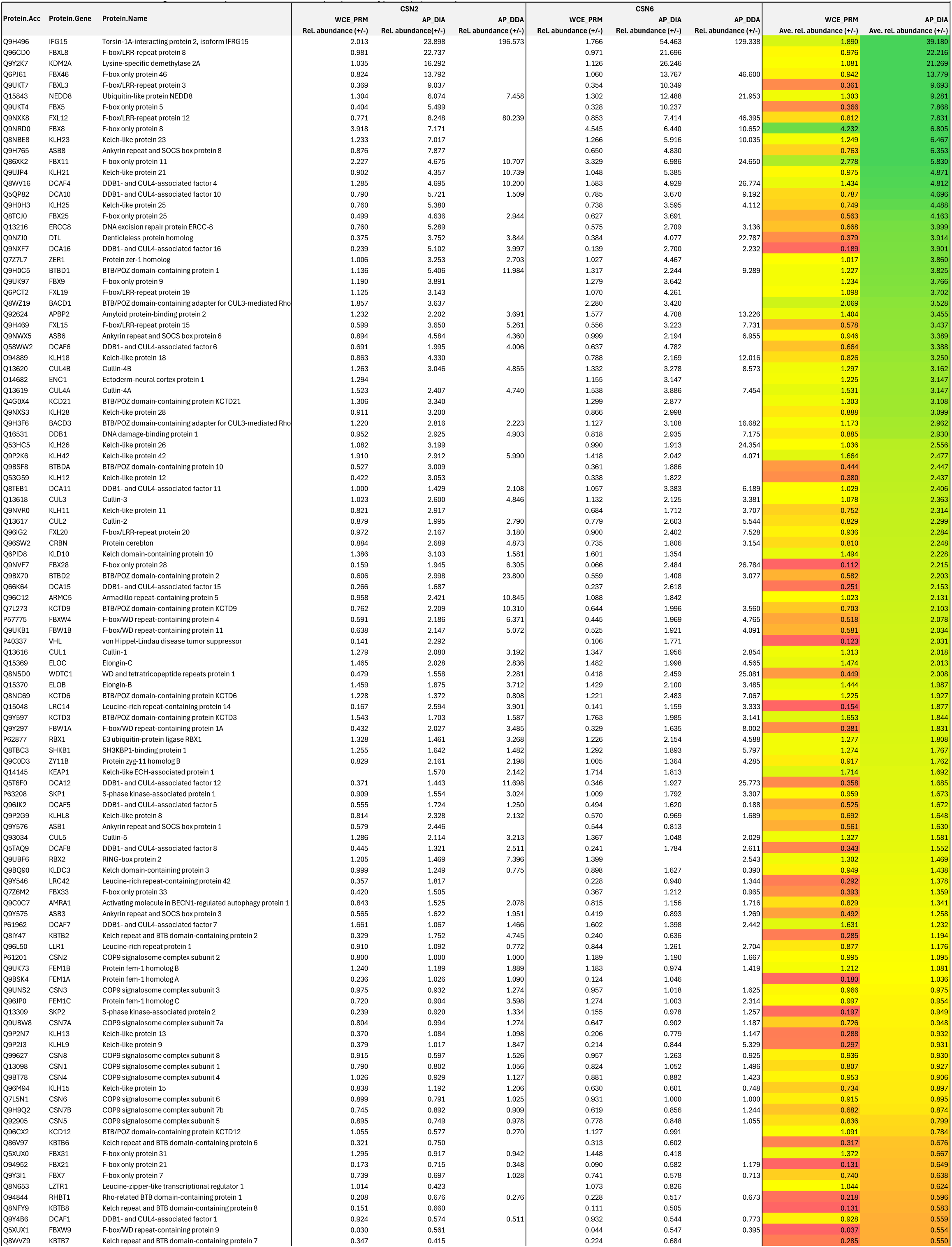
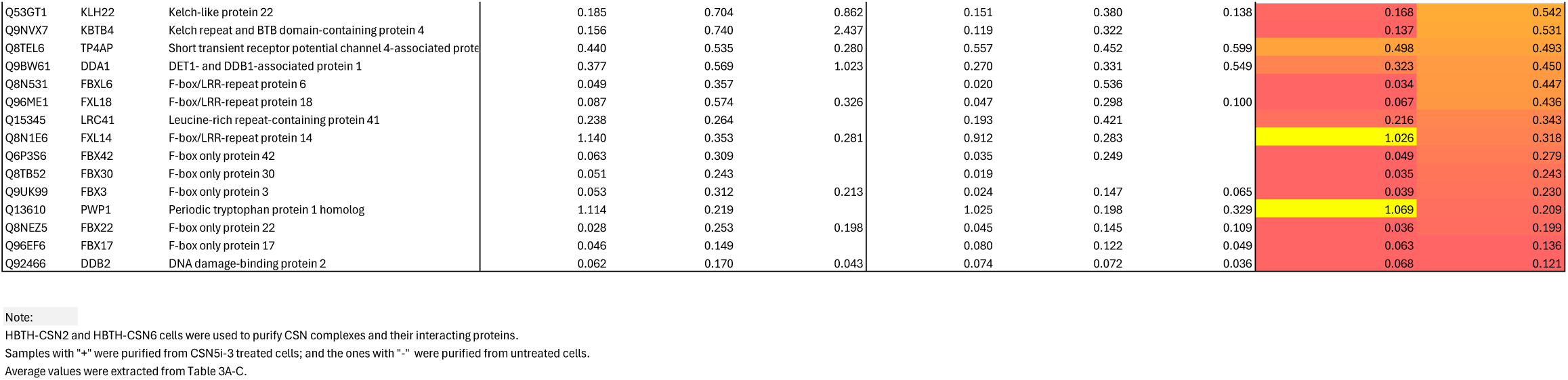
Relative abundance changes of the selected proteins in whole cell extracts (WCE) and affinity purified (AP) CSN complexes from CSN5i-3 treated and untreated cells.

## METHODS

### Protein purification and complex assembly

Seven of the eight human CSN subunits, excluding CSN5, were overexpressed and purified from *E. coli*. Two trimeric subcomplexes, CSN1–2–3 and CSN4–6–7, were individually purified through co-expression. For the CSN1–2–3 subcomplex, CSN2 was subcloned into a modified pGEX4T1 vector (Amersham Biosciences) containing an N-terminal glutathione S-transferase (GST) tag followed by a tobacco etch virus (TEV) protease cleavage site. Truncated CSN3 (residues 1–409) and full-length CSN1 were individually subcloned into a modified pET15b vector (Novagen) containing an N-terminal His tag with a TEV protease cleavage site. The expression cassettes of CSN1 and CSN3^1–409^, each driven by a T7 promoter and terminated by a T7 terminator, were subsequently inserted into the pGEX4T1-CSN2 plasmid. The resulting construct contained three independent expression cassettes for CSN2, CSN1, and CSN3^1–409^, respectively. The CSN1–2–3 complex was co-expressed in E. coli BL21(DE3) cells (Novagen), purified by glutathione-affinity chromatography, and subjected to TEV protease cleavage to remove the GST tag. The cleaved complex was further purified by anion exchange (Source Q, Cytiva) and size-exclusion chromatography (Superdex 200, Cytiva). The CSN4–6–7 subcomplex was prepared using the same strategy. In this case, truncated CSN7b (residues 1–239) was subcloned into the modified pGEX4T1 vector, while full-length CSN4 and CSN6 were inserted into the modified pET15b vector. CSN8 was subcloned into a modified pET15b vector encoding an N-terminal maltose-binding protein (MBP) tag followed by a TEV protease cleavage site. The overexpressed MBP–CSN8 fusion protein was initially purified by MBP affinity chromatography, followed by on-column TEV protease cleavage to remove the MBP tag. The cleaved protein was then further purified by ion-exchange chromatography. The CSN-7mer complex, comprising all subunits except CSN5, was reconstituted by incubating the CSN1–2–3, CSN4–6–7b, and CSN8 a molar ratio of 1:1.5:2. The assembled complex was subsequently purified by ion-exchange and size-exclusion chromatography.

CSN5 subunit was prepared from insect cells using a Bac-to-Bac expression system. Wild-type and mutant CSN5 were subcloned into a modified GTE vector (Invitrogen) containing an N-terminal octa-histidine (8×His) tag, followed by a Venus fluorescent protein and a TEV protease cleavage site. High-titer baculovirus was generated using ExpiSF9 insect cells (Thermo Fisher Scientific) cultured at 27 °C. For large-scale protein production, ExpiSF9 cells at a density of 2 × 10⁶ cells/mL were infected with baculovirus and cultured at 27 °C for 68 hours. Cells were harvested by centrifugation, resuspended in lysis buffer containing 20 mM Tris-HCl pH 8.0, 200 mM NaCl, 1 mM PMSF, and Roche Complete Protease Inhibitor, and lysed by sonication. The lysate was clarified by centrifugation, and CSN5 was purified through a series of chromatography steps, including His-tag affinity, ion-exchange, and size-exclusion chromatography. To obtain untagged CSN5, the Venus tag was removed by TEV protease digestion performed during dialysis following the initial His-affinity purification. The resulting mixture was subsequently passed through a second Ni-NTA column to separate untagged CSN5 from the cleaved His–Venus tag. To assemble the complete CSN complex, purified CSN5 was incubated with tag-free CSN-7mer at a 2:1 molar ratio on ice for 30 minutes, then purified by size-exclusion chromatography (Superdex 200) using a buffer containing 20 mM HEPES (pH 7.5), 150 mM NaCl, and 0.2 mM TCEP. The peak fractions containing CSN complex were pooled, concentrated, flash-frozen, and stored at −80°C.

### Preparation of N8∼CRL1 protein complexes

Two short unstructured segments in the N-terminus of CUL1 (residues 1–12 and 58–81) were removed from the full-length human CUL1, resulting in CUL1^ΔN^ (referred to here as CUL1). Both CUL1 and RBX1 (residues 16–108) were fused with an N-terminal 6xHis tag followed by a TEV cleavage site and co-expressed in BL-21(DE3). The complex was initially purified using Ni²⁺-Sepharose affinity chromatography. Following TEV cleavage to remove the His tag, the complex was further purified by cation-exchange and size-exclusion chromatography. The same strategy was used to prepare other CRL complexes, including CUL2-RBX1, CUL3-RBX1, CUL4A-RBX1 and CUL5-RBX2.

To prepare the heterodimeric NEDD8-activating enzyme APPBP1–UBA3, APPBP1 was subcloned into the modified pGEX4T1 vector containing a GST tag followed by a TEV protease cleavage site, while UBA3 was subcloned into a modified pET15b vector containing a chloramphenicol resistance cassette. GST–APPBP1 and UBA3 were co-expressed in *E. coli* BL21(DE3) and purified by glutathione-affinity chromatography. After TEV cleavage, the APPBP1–UBA3 complex was further purified by anion-exchange and size-exclusion chromatography. The NEDD8-conjugating enzyme UBC12 and NEDD8 (wild-type and mutant forms) were subcloned into the same modified pGEX4T1 vector, expressed in *E. coli* BL21(DE3), and purified via glutathione-affinity and anion-exchange chromatography. A truncated form of NEDD8 ending at glycine 76, representing the mature processed form, was used throughout this study. To generate the neddylated CRL1 complex (N8∼CRL1), 10 μM of purified CRL1 was incubated with 10 μM GST–NEDD8 in the presence of 0.2 μM APPBP1–UBA3 and 0.5 μM UBC12 at 4 °C for 1 hour. The neddylation reaction was carried out in buffer containing 20 mM Tris-HCl pH 8.0, 200 mM NaCl, 0.4 mM TCEP, 2 mM ATP and 10 mM MgCl_2_. The resulting GST-N8∼CRL1 was separated from unmodified CRL1 by glutathione-affinity chromatography. After on-column TEV cleavage, N8∼CRL1 was eluted from the column and further purified by cation-exchange and gel filtration chromatography.

To conveniently monitor the production of free NEDD8 during CSN-mediated deneddylation, NEDD8 was site-specifically labeled with the fluorescent dye Alexa Fluor 633 (Sigma) via a cysteine residue engineered in place of the native N-terminal methionine. Excess dye was removed by size-exclusion chromatography. The fluorescently labeled NEDD8 was then conjugated to the CUL–RBX1 complex using the neddylation protocol described above. For BLI experiments measuring the binding of NEDD8 to CSN, an AviTag followed by a biologically inert GB1 tag (the B1 domain of streptococcal protein G; 56 residues, ∼7 kDa) was fused to the N-terminus of NEDD8. The purified Avi–GB1–NEDD8 fusion protein was efficiently biotinylated in vitro using *E. coli* biotin ligase (BirA). Excess free biotin was removed by size-exclusion chromatography on a Superdex 75 column, yielding monodisperse, biotinylated Avi–GB1–NEDD8 suitable for immobilization on streptavidin biosensors.

### BioLayer interferometry

The binding affinity between NEDD8 and CSN or CSN5–CSN6 was measured using the Octet Red 96 system (Sartorius). SA biosensors (Sartorius) coated with streptavidin were loaded with 200 nM biotinylated NEDD8 and subsequently quenched with 200 nM biocytin before performing binding analyses. The reactions were conducted in black 96-well plates maintained at 30°C. The binding buffer contained 20 mM HEPES pH 7.5, 150 mM NaCl, 0.2 mM TCEP, 0.1% Tween-20, and 0.1 mg/mL ovalbumin. CSN or CSN5–CSN6 were tested as analytes at 3-fold serially diluted concentrations in the absence or presence of 20 µM CSN5i-3. To assess the binding affinity between CSN^DM^ and N8∼CRL1 containing wild type or 3A and 3S mutant RBX1, biotinylated N8∼CRL1 was initially loaded to streptavidin-coated biosensors. CSN^DM^ was tested as analyte at 3-fold serially diluted concentrations. To measure the binding affinity of CSN5i-3 with CSN or CSN5–CSN6, biotinylated anti-Venus nanobody was initially bound to streptavidin-coated biosensors to convert the probes to Venus-nanobody biosensors. Venus-tagged CSN was then loaded through the interaction between Venus and the anti-Venus nanobody. To measure the binding of CSN5i-3 to CSN5–CSN6, biotinylated CSN5–CSN6 protein was loaded to streptavidin-coated biosensors. CSN5i-3 were measured as analyte at 3-fold serially diluted concentrations. Data analysis of all BLI experiments was performed using Octet data analysis software, and the dissociation constant (K_D_) was determined from either kinetics or steady-state equilibrium measurements. All BLI experiments were performed a minimum of three times.

### Isothermal titration calorimetry

Isothermal titration calorimetry (ITC) (Marven) binding assays were performed at 25°C using a MicroCal PEAQ-ITC system (Malvern). To measure the binding of CSN5i-3 or CSN5i-1a to CSN5–CSN6, 160 µM CSN5–CSN6 in the syringe was titrated into 10 µM CSN5i-3 or CSN5i-1a in the cell. The data were fitted to a one-site binding model to determine the dissociation constants (K_D_), enthalpy (ΔH), and stoichiometry (N) using the MicroCal PEAQ-ITC Analysis Software.

### Cryo-EM sample preparation and data collection

For grid preparation of all CSN complexes, holey gold grids (UltraAuFoil R1.2/1.3, 300 mesh) were pretreated with glow discharge. FOM was added to all samples at a final concentration of 0.07% (w/v) before grid preparation. For CSN5i-3-CSN, CSN5i-3-CSN-NEDD8, CSN5i-3-CSN-N8∼CRL1, and CSN5i-1a-CSN complexes, 10 μM CSN was incubated first with 50 μM CSN5i-3 or CSN5i-1a and then with 30 μM NEDD8 or N8∼CRL1. Three microliters of the sample were applied to grid. For the CSN^DM^-N8∼CRL (CRL1, CRL2, CRL3, CRL4A, and CRL5) complexes, N8∼CRLs were mixed with CSN^DM^ at a 3:1 ratio and incubated for 30 minutes on ice. The concentration was diluted to approximately 4.5 mg/ml for grid preparation. The grids were subsequently blotted for 6 s (blotting force at 0, temperature at 10 ℃, and relative humidity at 100%), plunged and flash frozen into liquid ethane using a Vitrobot Mark IV, and stored in liquid nitrogen for data collection.

Multiple complexes were imaged at different transmission electron microscopes at different Cryo-EM facilities. For the CSN, CSN^DM^-N8∼CRL4, CSN5i-3-CSN-NEDD8 and CSN5i-1a-CSN complexes, data collection was carried out on a Titan Krios transmission electron microscope (Thermo Fisher Scientific) operated at 300 kV at the University of Washington. For the CSN^DM^-N8∼CRL1, CSN^DM^-N8∼CRL2, CSN5i-3-CSN-N8∼CRL1 complex, data collection was carried out on a Titan Krios transmission electron microscope operated at 300 kV at the HHMI Janelia Research Campus. For CSN^DM^-N8∼CRL5 complex, data collection was carried out on a Titan Krios transmission electron microscope operated at 300 kV at the National Cancer Institute. The automation scheme was implemented using the SerialEM software^41,42^ with beam-image shift strategy and active beam-tilt compensation at a nominal magnification of 105,000x, resulting in a physical pixel size of 0.840 Å, 0.827 Å, 0.835 Å, respectively. The zero-loss-energy images were acquired on a Gatan K3 direct detector operated in correlated double sampling mode with the slit width of post-column Gatan BioQuantum GIF energy filter set to be 20, 20, 10-eV, respectively. The dose rate was adjusted to 13.9, 10.1, 14.6 e^−^ per Å^2^ per second, respectively, and a total dose of 60 e^−^ per Å^2^ for each image fractionated into 60 frames. The images were recorded at a defocus range of −0.8∼3.0 μm. For the CSN^DM^-N8∼CRL3 and CSN5i-3-CSN complexes, data collection was carried out on a 200 kV Talos Glacios transmission electron microscope at University of Washington. The data collection performed with SerialEM automation scheme via beam-image shift strategy at a nominal magnification of 45,000x, resulting in a physical pixel size of 0.885 Å, using Gatan K3 direct detector operated in correlated double sampling mode. The dose rate was adjusted to 17.3 e^−^ per Å^2^ per second, and a total dose of 50 e^−^ per Å^2^ for each image fractionated into 100 frames. The images were recorded at a defocus range of −0.8∼2.5 μm.

### Cryo-EM data processing

All datasets were processed with cryoSPARC^43^. The beam-induced motion correction and CTF estimation was conducted with cryoSPARC Live session using Patch Motion Correction and patch CTF estimation. The motion-corrected micrographs were filtered by defocus, CTF fit resolution (excluding >8 Å), and relative ice thickness. The curated micrographs were exported and subject to blob picking. After inspection of picked particles and removal of targets on ice or Au foil, the left particles were extracted with fourfold binning and subject to 2D classification. The good particles were kept and subject to ab initio reconstruction with 4 classes. All particles then subject to heterogeneous refinement with the four reconstructions generated from ab initio reconstruction as reference. Approximately 100,000 particles were randomly selected after heterogenous refinement. Particles of good reconstructions were used as template to train model for Topaz picking^44^. After picked by topaz, particles were extracted and cleaned with 2D classification. Good particles were kept and filtered by center-to-center distance to remove duplicate particles.

For CSN5i-3-CSN-N8∼CRL1, 229,370 good particles from 7,564 micrographs generated from sequential blob picking, 2D classification were subjected to sequential ab-initio reconstruction and heterogenous refinement. 149, 404 particles from good reconstruction were used for topaz training. 1,174,702 particles were picked using topaz picking. After one round 2D classification and duplicates curation, 536,325 particles were kept and subjected to heterogeneous refinement using the four reconstructions generated from previous ab-initio reconstruction as reference. 370,244 particles were kept and subjected to sequential non-uniform refinement and CTF refinement. These particles yield a 2.9 Å reconstruction. 3D classification was applied to further improve the density of CSN5-CSN6. 174, 577 particles were kept and yield a 3.0 Å reconstruction. The following local refinement with a binary mask focused on CSN5-CSN6-N8∼CUL1-WHB domain yields a 3.3 Å reconstruction.

For CSN^DM^-N8∼CRL1, 726,752 good particles from 10,052 micrographs were subjected to heterogenous refinement. 434,896 particles from one good class at 5.2 Å were kept for sequential particle extraction and non-uniform refinement^45^. After refinement, 433,043 particles yielded a nominal 3.3 Å reconstruction. The densities of CSN5-CSN6 and N8∼CRL1 were less resolved than the C-terminal helical bundle of CSN. To further improve the density of the catalytic hemisphere, the particles were subject to 3D variability analysis with focused mask containing CSN5-CSN6 globular domain, NEDD8, WHB, and RBX1 globular domains implemented. After 3D variability analysis^46^, 177,066 particles of improved density covering CSN5-CSN6 and N8∼CUL1-WHB were kept, which yielded a 3.2 Å reconstruction. After sequential global CTF, local refinement and non-uniform refinement, a 3.2 Å reconstruction was obtained. To further polish the CSN5-CSN6-NEDD8 density, same binary mask as 3D Variability imposed for local refinement. The local refinement improved the CSN5-CSN6 density and allowed us to visualize the sidechain density of NEDD8, CSN5-CSN6, and the CUL1WHB domain. More detailed information could be found in Extended Data Fig. 1, 3, and 4, as well as Extended Data Table 1.

### Model building and refinement

For the CSN5i-3-CSN structure, the PDB model 4D10 was fitted into the 3.3 Å CSN-CSN5i-3 EM map using Chimera^47^. The fitted model was used as the initial structure to rebuild the CSN complex. The CSN7a from 4D10 served as a template to trace the main chain, and the residue sequence was modified to match CSN7b in Coot^48^. For the flexible regions of CSN2 and CSN4, the PDB model was trimmed based on the density in Coot. The CSN5i-3 model from 5JOG was fitted into the 3.3 Å CSN5i-3-CSN EM map and refined using Phenix. For the CSN5i-1a-CSN structure, CSN5i-1b in PDB model 5JOH was modified based on CSN5i-1a SMILES file. Then CSN model from CSN5i-3-CSN and CSN5i-1a was fitted into the 3.0 Å CSN5i-1a-CSN EM map and refined with Phenix. For the CSN5i-3-CSN-N8∼CRL1 model, the previous CSN-CSN5i-3 structure was used as the starting model. The CSN5-CSN6, CSN2, and CSN4 subunits undergo conformational changes upon binding to N8∼CRL1. Their structural models were first split and then fitted into the 3.3 Å EM map. Residues were adjusted in Coot based on sequence and side-chain density. The PDB model 1LDJ of CRL1 was fitted and manually adjusted in Coot. The iso-peptide bond was manually built in UCSF ChimeraX. For the CSN5i-3-CSN5-NEDD8 structure, CSN5i-3-CSN and NEDD8 from the PDB model 1XT9 were fitted into the EM maps and adjusted in Coot. The well-resolved density allowed us to model CSN5i-3 and the zinc ions. For the other CSN^DM^-N8∼CRL complexes, the CSN5i-3-CSN-N8∼CRL1 model was fitted into the EM map and adjusted in Coot. For the rest of the CRLs, the N8∼CRL structures were automatically built using ModelAngelo^49^, combined with the docked CSN^DM^ model then further refined manually in Coot. The final model was subjected to real space refinement in Phenix^50^. The iso-peptide bond between the NEDD8 C-terminal glycine residue and the neddylation site lysine of CRLs was manually built and refined with Phenix. The structural figure panels were prepared with PyMol and UCSF ChimeraX.

### DSSO cross-linking of the CSN-N8∼CR1 complex

To assemble the CSN-N8∼CRL1 complex *in vitro* for cross-linking, 7 µL of the purified CSN (6.8 μM) and 4.8 μL of N8∼CRL1 (30 μM) complexes were mixed and incubated on ice for 10 s or 30 min. For the treated samples, CSN was first incubated with CSN5i-3 at a molar ratio of 1:10 for 30 min on ice before mixing with N8∼CRL1. After incubation, the complexes were cross-linked with DSSO at a molar ratio of 1:100 (protein to linker) for 1 hr at room temperature^51^. After quenching with 50 mM NH_4_HCO_3_ for 15 min, cross-linked CSN-N8∼CRL1 complexes were transferred onto a 30 KDa microcon and washed with 25 mM NH_4_HCO_3_. The proteins were reduced by 3.3 mM TCEP for 30 min and alkylated with 16.5 mM iodoacetamide for 30 min in dark. After washing with 25 mM NH_4_HCO_3_, proteins were reconstituted in 60 µL of 8 M urea in 25 mM NH_4_HCO_3_. Lys-C (enzyme to protein ratio of 1:100) was added to the solution and the mixture was incubated at 37°C for 4 hr. Then, urea was reduced to 1.5 M for trypsin digestion (enzyme to protein ration of 1:50) at 37°C overnight. The peptide digests were extracted, desalted by Sep-Pak C18 cartridge (Waters) and stored in −80°C before TMT labeling.

### TMT labeling and SEC-HpHt fractionation

Equal amounts of cross-linked peptides from each condition were labeled with 10-plex TMT reagents (Thermo Fisher Scientific) according to manufacturer’s protocol. Briefly, peptides (14.5 μg) were dissolved in 25 μL of 100 mM TEAB (pH 8.5). 0.8mg of TMT reagents were dissolved in 41 μL of anhydrous acetonitrile (Sigma), and 3 μL of each TMT reagent was added to the corresponding aliquot of peptides together with 7 μL of anhydrous ACN. The reaction was incubated for 1 hr with shaking and quenched with 5% hydroxylamine for 15 min at room temperature. The labeled peptides were pooled and dried using Speed-Vac. The dried samples were resuspended in 0.1% TFA buffer and desalted using Sep-Pak C18 cartridge and dried. The TMT labeled peptides were dissolved in 30% ACN/0.1% TFA, mixed and fractionated on a Superdex™ 30 Increase 3.2/300 column with an Agilent 1260 Series HPLC. The two fractions containing inter-linked peptides (F23-25min, F25-27min) were collected for direct LC-MS^n^ analysis or further HpHt separation^52^. For HpHt separation, each SEC fraction was dissolved in 160 µL of ammonia water (pH 10) and loaded onto a HpHt tip. Peptides were eluted with increasing percentage of ACN in ammonia water (6%, 9%, 12%, 15%, 18%, 21%, 25%, 30%, 35%, and 50%). The 25%, 30%, 35% and 50% fractions were combined to 6%, 9%, 12% and 21% fractions, respectively for LC MS^n^ analysis (Jiao, 2022).

### LC MS^n^ analysis

Cross-linked peptides were analyzed by LC MS^n^ using an UltiMate 3000 RSLC coupled with an Orbitrap Fusion Lumos mass spectrometer. Samples were loaded onto a 50 cm x 75 μm Acclaim PepMap C18 column and separated over a 180 min gradient of 4% to 25% acetonitrile at a flow rate of 300 nL/min (Jiao, 2022). Ions with charge of 4+ to 8+ in the MS^1^ scan were selected for MS^2^ analysis. The top 4 most abundant fragment ions in MS^2^ scan were selected for MS^3^ sequencing. The CID-MS^2^ normalized collision energy was 23%. For MS^3^ scans, CID was used with a collision energy of 35%. TMT quantitation on cross-linked peptides were accomplished using the Synchronous Precursor Selection (SPS) MS^3^ method^28^.

### Identification and quantitation of cross-linked peptides

MS^3^ spectra was extracted by PAVA (UCSF) and subjected to Protein Prospector (v.6.3.5) for database searching (using Batch-Tag against SwissProt human database, version 2024. 08. 05, 20,436 entries). The mass tolerances were set as ± 20 ppm for parent ions and 0.6 Da for fragment ions. Trypsin was set as the enzyme with three maximum missed cleavages allowed. Cysteine carbamidomethylation and TMTplex at protein N-terminus were set as static modifications. A maximum of four variable modifications were also allowed, including TMTplex at lysine, methionine oxidation, N-terminal acetylation, and N-terminal conversion of glutamine to pyroglutamic acid. Three defined DSSO cross-linked modification on uncleaved lysines, including alkene (C3H2O, +54 Da), thiol (C3H2SO, +86 Da) and sulfenic acid (C3H4O2S, +104Da) were also selected as variable modifications. The in-house software XL-Tools was used to automatically identify, summarize and validate cross-linked peptides based on Protein Prospector database search results and MS^n^ data^51^. Quantification values for cross-linked peptides were obtained from reporter ion intensities in SPS MS^3^ scans^28^. These intensities were corrected for isotopic impurities of the different TMTplex reagents based on the manufacturer’s specifications.

### Affinity purification of CSN complexes

HEK293T cells expressing HBTH-tagged CSN2 and CSN6 were grown to ∼90% confluence in DMEM medium containing 10% FBS and 1% Pen/strep^32^. The cells were treated with 1μM CSN5i-3 or DMSO for 16 hr. Next, in-cell cross-linking was performed with 0.025% formaldehyde for 10 min at 37 °C in PBS buffer^31^. The cells were then washed with PBS and lysed in a native lysis buffer containing 150 mM sodium chloride, 50 mM sodium phosphate, 10% glycerol, 1 mM ATP, 1 mM DTT, 5 mM MgCl_2_, 1X protease inhibiter, 1X phosphatase inhibitor, and 0.5% NP-40, pH 7.5. The lysates were centrifuged at 13,000 rpm for 15 min to remove cell debris, and the supernatant was incubated with streptavidin resin for 2 hr at 4°C. The bead-bound proteins were washed with 50 volumes of lysis buffer followed by 20 volumes of another buffer containing150 mM NaCl, 25 mM Na_2_HPO_4_, and 5% glycerol, pH 7.6, and then cross-linked with 0.5 mM DSSO for 1 hr at 37°C. The cross-linked CSN complexes were reduced and alkylated then digested with LyC/Trypsin. The digested peptides were desalted with C18 tips prior to liquid chromatography tandem mass spectrometry (LC-MS/MS).

### Digestion of cell lysates

To examine protein abundance changes in cells during CSN5i-3 treatment, we digested 1mg cell lysates from treated and untreated cells using a FASP protocol (Jiao, 2022). Proteins were reduced, alkylated and digested on 30 KD microcon (Millipore). The digested peptides were desalted with Sep-Pak column prior to LC MS/MS.

### LC MS/MS analysis for protein identification and label-free quantitation

For all LC-MS acquisitions, HPLC separation was performed on an UltiMate 3000 UHPLC (Thermo Fisher Scientific) using an EasySpray 50cm x 75μm I.D. Acclaim® PepMap RSLC column heated at 45° C and coupled on-line to an Orbitrap Fusion Lumos mass spectrometer (Thermo Fisher Scientific). Specifically, a 4-22% Buffer B gradient (Buffer A: 0.1% formic acid; Buffer B: 100% acetonitrile containing 0.1% formic acid) was run either over 57 minutes (1.5-hour method) or 87 minutes (2-hour method).

### DDA-based analysis

The DDA-MS method for affinity-purified CSN complexes was 90 min and consisted of an MS^1^ survey scan (375–1800 m/z, resolution of 120,000 at m/z 200, AGC target 4.0 × 10^5^, maximum injection time 50 ms) followed by data-dependent MS/MS acquired in the linear ion trap with HCD NCE30 at top speed for 3 sec. Target ions already selected for MS/MS were dynamically excluded for 30 sec. MS^2^ scans were acquired in the linear ion trap using ‘Rapid’ mode with AGC target 2.0 × 10^4^ with a maximum injection time of 35 ms. Protein quantitation was carried out using MaxQuant (v2.0.3.0) against human proteome sequences from SwissProt database (20,418 entries; March 2025 version)^53^. The first search peptide tolerance was set to 15 ppm, with main search peptide tolerance set to 4.5 ppm. Both peptide spectrum match and protein false discovery rates (FDRs) were set at 1%, in razor peptide fashion. Trypsin was selected for the protease with up to 1 missed cleavage; no nonspecific cleavage was allowed. For protein quantitation, cysteine carbamidomethylation was set as a fixed modification, while methionine oxidation and N-terminal acetylation were selected for variable modifications, maximum of 1 per peptide. Intensities were determined as the full peak volume over the retention time profile. Intensities of different isotopic peaks in an isotope pattern were always summed up for further analysis. “Unique plus razor peptides” was selected as the degree of uniqueness required for peptides to be included in quantification.

### Data-independent acquisition (DIA)-based analysis

The DIA-MS method for affinity-purified CSN complexes was 90 min and consisted of an MS^1^ survey scan followed by 33 MS^2^ scans using variable windows^54^. The MS^1^ scan range was set as 350–1650 m/z with a resolution of 120,000 at m/z 200. The AGC target value for the MS^1^ scan was set to 2.0 × 10⁶ with a maximum injection time of 50 ms. MS^2^ scans were acquired at a resolution of 30,000 (at m/z 200), using HCD NCE30. For all DIA analysis, including library generation, protein identification and quantitation, DIA-NN (v2.0.2) was used^55^. The spectral library was generated using human proteome sequences from SwissProt database (20,418 entries; March 2025 version). Trypsin was set as the protease with 1 missed cleavage. N-term excision and cysteine carbamidomethylation were set as fixed modifications, while methionine oxidation was set as a variable modification with 1 max per peptide. The peptide length was set as 7∼30 aa, with charge 2∼4. Precursor FDR was set to 1% for the output and proteins were quantified using the QuantUMS strategy^56^.

### Parallel reaction monitoring (PRM)-based targeted quantitative analysis

To generate the PRM target library for quantitation, peptide digests from affinity-purified CSN complexes were pooled and run using 2 hr DIA-MS methods with the same settings as outlined above. The top 3 abundant peptide ions from the selected 126 proteins were chosen, with preference towards unmodified peptides and sequences identified without missed cleavages where possible. A total of 351 precursors from 126 proteins were selected and monitored across two 2 hr acquisitions for cell lysate samples, with each peptide being scheduled within a 4 min window. The isolation window was set as 0.7 Da and Orbitrap resolution was 30,000 at m/z 200. Transitions determined by DIA-MS analysis were quantified using Skyline (v23.1.0.268)^57^. The peak areas of all transitions were summed for each peptide and then averaged to represent protein abundances. Final protein abundances were then averaged across biological replicates and compared between treated and control lysates. Two biological replicates were performed for each condition.

### Validation of the selected CSN interactions by immunoblotting analysis

To validate the selected CSN interactions, affinity purified human CSN complexes were separated by SDS−PAGE. Proteins were transferred to a PVDF membrane and analyzed by immunoblotting. HBTH-CSN2 and HBTH-CSN6 were detected using a streptavidin−HRP conjugate (1:5000), DDB2 was detected by using anti-DDB2 antibody (life technology MA5-34832), Cul4A was detected by using anti-Cul4A (life technology PA5-14542), and FBX022 was detected by using anti-FBX022 (protein tech 13606-1-AP).

### Affinity pull-down analysis

Approximately 200 µg of purified CSN complex with His–Venus–tagged CSN5 was used as the bait. Venus–tagged NEDD8 (wild-type and three mutants) were added at a 25:1 molar excess relative to CSN, either in the presence or absence of 10 µM CSN5i-3. The bait and binding partners were incubated together on ice at 4 °C for 2 h with gentle mixing. The reaction mixtures were then applied to 50 µL of Ni–NTA Sepharose resin and incubated with end-over-end rotation at 4 °C for 3 h. Beads were washed extensively with a binding buffer containing 20 mM Tris-HCl, pH 8.0, 150 mM NaCl, and 10 mM imidazole. Bound complexes were eluted in 100 µL of an elution buffer containing 20 mM Tris-HCl, pH 8.0, 150 mM NaCl, and 300 mM imidazole. Eluates were resolved by SDS–PAGE and the Venus signal was detected by in-gel fluorescence imaging.

### Inhibition of iso-peptidase activity by CSN5i-3 and CSN5i-1a

The enzymatic activities of CSN and CSN5-CSN6 were determined by monitoring the amounts of the deneddylation product, NEDD8 conjugated with the fluorescent dye CF633, and the reaction substrate, CF633-N8∼CRL1. Twenty-microliter reactions were carried out in PCR tubes. Enzymes (recombinant CSN or CSN5-CSN6) were used at 2.0 nM (added at T=0), and CF633-N8∼CRL1 was used at 2 µM. CSN5i-3 was serially diluted 3-fold from 450 nM down to 0.062 nM. CSN5i-1a was serially diluted 3-fold from 100 µM down to 137 nM. Reactions at room temperature were stopped after 25 minutes by adding 8 µL of 4X SDS loading buffer containing Orange G, then heated for 5 minutes at 90°C. Two tubes without inhibitor were designated as 100% activity, and one tube that contained SDS loading buffer prior to enzyme addition was considered 0% activity. The reaction products NEDD8-CF633 and the CF633-N8∼CRL1 substrate were separated on 4–15% mini-Protein gels (BioRad) at 250 V (67 mA) for 20–30 minutes. Fluorescence signals from CF633 were detected using an Odyssey CLX Imaging System (LICOR) in the 700 nm channel. All fluorescence signals were normalized to correct for loading variations. The Michaelis constant (Km) was determined for both CSN and CSN5-CSN6 with the CF633-N8∼CRL1 substrate under the same buffer and reaction conditions. The substrate was serially diluted 3-fold from 1.0 µM to 0.021 nM. Fluorescence signals from CF633 were detected using the Odyssey CLX Imaging System (LICOR) in the 700 nm channel, and the enzyme rate versus substrate concentration was analyzed using an enzymatic kinetic model in Prism (GraphPad).

## REFERENCES

1. Copeland, R.A. (2013). Why Enzymes as Drug Targets? Enzymes Are Essential for Life. In Evaluation of Enzyme Inhibitors in Drug Discovery: A Guide for Medicinal Chemists and Pharmacologists, (Wiley-Interscience), pp. 1–33. 10.1002/9781118540398.

2. Lin, H. (2023). Substrate-selective small-molecule modulators of enzymes: Mechanisms and opportunities. Curr Opin Chem Biol 72, 102231. 10.1016/j.cbpa.2022.102231.

3. Qin, N., Xu, D., Li, J., and Deng, X.W. (2020). COP9 signalosome: Discovery, conservation, activity, and function. J Integr Plant Biol 62, 90–103. 10.1111/jipb.12903.

4. Schulze-Niemand, E., and Naumann, M. (2023). The COP9 signalosome: A versatile regulatory hub of Cullin-RING ligases. Trends Biochem Sci 48, 82–95. 10.1016/j.tibs.2022.08.003.

5. Singh, A.K., and Chamovitz, D.A. (2019). Role of Cop9 Signalosome Subunits in the Environmental and Hormonal Balance of Plant. Biomolecules 9. 10.3390/biom9060224.

6. Harper, J.W., and Schulman, B.A. (2021). Cullin-RING Ubiquitin Ligase Regulatory Circuits: A Quarter Century Beyond the F-Box Hypothesis. Annu Rev Biochem 90, 403–429. 10.1146/annurev-biochem-090120-013613.

7. Cope, G.A., and Deshaies, R.J. (2003). COP9 signalosome: a multifunctional regulator of SCF and other cullin-based ubiquitin ligases. Cell 114, 663–671.

8. Rao, F., Lin, H., and Su, Y. (2020). Cullin-RING Ligase Regulation by the COP9 Signalosome: Structural Mechanisms and New Physiologic Players. Adv Exp Med Biol 1217, 47–60. 10.1007/978-981-15-1025-0_4.

9. Mayor-Ruiz, C., Jaeger, M.G., Bauer, S., Brand, M., Sin, C., Hanzl, A., Mueller, A.C., Menche, J., and Winter, G.E. (2019). Plasticity of the Cullin-RING Ligase Repertoire Shapes Sensitivity to Ligand-Induced Protein Degradation. Mol Cell 75, 849–858.e848. 10.1016/j.molcel.2019.07.013.

10. Xue, G., Xie, J., Hinterndorfer, M., Cigler, M., Dötsch, L., Imrichova, H., Lampe, P., Cheng, X., Adariani, S.R., Winter, G.E., and Waldmann, H. (2023). Discovery of a Drug-like, Natural Product-Inspired DCAF11 Ligand Chemotype. Nat Commun 14, 7908. 10.1038/s41467-023-43657-6.

11. Hsia, O., Hinterndorfer, M., Cowan, A.D., Iso, K., Ishida, T., Sundaramoorthy, R., Nakasone, M.A., Imrichova, H., Schätz, C., Rukavina, A., et al. (2024). Targeted protein degradation via intramolecular bivalent glues. Nature 627, 204–211. 10.1038/s41586-024-07089-6.

12. Mayor-Ruiz, C., Bauer, S., Brand, M., Kozicka, Z., Siklos, M., Imrichova, H., Kaltheuner, I.H., Hahn, E., Seiler, K., Koren, A., et al. (2020). Rational discovery of molecular glue degraders via scalable chemical profiling. Nat Chem Biol 16, 1199–1207. 10.1038/s41589-020-0594-x.

13. Wang, K., Deshaies, R.J., and Liu, X. (2020). Assembly and Regulation of CRL Ubiquitin Ligases. Adv Exp Med Biol 1217, 33–46. 10.1007/978-981-15-1025-0_3.

14. Lyapina, S., Cope, G., Shevchenko, A., Serino, G., Tsuge, T., Zhou, C., Wolf, D.A., Wei, N., and Deshaies, R.J. (2001). Promotion of NEDD-CUL1 conjugate cleavage by COP9 signalosome. Science 292, 1382–1385. 10.1126/science.1059780.

15. Kurz, T., Ozlü, N., Rudolf, F., O’Rourke, S.M., Luke, B., Hofmann, K., Hyman, A.A., Bowerman, B., and Peter, M. (2005). The conserved protein DCN-1/Dcn1p is required for cullin neddylation in C. elegans and S. cerevisiae. Nature 435, 1257–1261. 10.1038/nature03662.

16. Cope, G.A., Suh, G.S., Aravind, L., Schwarz, S.E., Zipursky, S.L., Koonin, E.V., and Deshaies, R.J. (2002). Role of predicted metalloprotease motif of Jab1/Csn5 in cleavage of Nedd8 from Cul1. Science 298, 608–611. 10.1126/science.1075901.

17. Lingaraju, G.M., Bunker, R.D., Cavadini, S., Hess, D., Hassiepen, U., Renatus, M., Fischer, E.S., and Thomä, N.H. (2014). Crystal structure of the human COP9 signalosome. Nature 512, 161–165. 10.1038/nature13566.

18. Cavadini, S., Fischer, E.S., Bunker, R.D., Potenza, A., Lingaraju, G.M., Goldie, K.N., Mohamed, W.I., Faty, M., Petzold, G., Beckwith, R.E., et al. (2016). Cullin-RING ubiquitin E3 ligase regulation by the COP9 signalosome. Nature 531, 598–603. 10.1038/nature17416.

19. Faull, S.V., Lau, A.M.C., Martens, C., Ahdash, Z., Hansen, K., Yebenes, H., Schmidt, C., Beuron, F., Cronin, N.B., Morris, E.P., and Politis, A. (2019). Structural basis of Cullin 2 RING E3 ligase regulation by the COP9 signalosome. Nat Commun 10, 3814. 10.1038/s41467-019-11772-y.

20. Enchev, R.I., Scott, D.C., da Fonseca, P.C., Schreiber, A., Monda, J.K., Schulman, B.A., Peter, M., and Morris, E.P. (2012). Structural basis for a reciprocal regulation between SCF and CSN. Cell Rep 2, 616–627. 10.1016/j.celrep.2012.08.019.

21. Mosadeghi, R., Reichermeier, K.M., Winkler, M., Schreiber, A., Reitsma, J.M., Zhang, Y., Stengel, F., Cao, J., Kim, M., Sweredoski, M.J., et al. (2016). Structural and kinetic analysis of the COP9-Signalosome activation and the cullin-RING ubiquitin ligase deneddylation cycle. Elife 5. 10.7554/eLife.12102.

22. Hu, Y., Zhang, Z., Mao, Q., Zhang, X., Hao, A., Xun, Y., Wang, Y., Han, L., Zhan, W., Liu, Q., et al. (2024). Dynamic molecular architecture and substrate recruitment of cullin3-RING E3 ligase CRL3. Nat Struct Mol Biol 31, 336–350. 10.1038/s41594-023-01182-6.

23. Schlierf, A., Altmann, E., Quancard, J., Jefferson, A.B., Assenberg, R., Renatus, M., Jones, M., Hassiepen, U., Schaefer, M., Kiffe, M., et al. (2016). Targeted inhibition of the COP9 signalosome for treatment of cancer. Nat Commun 7, 13166. 10.1038/ncomms13166.

24. Zhang, Y., Jost, M., Pak, R.A., Lu, D., Li, J., Lomenick, B., Garbis, S.D., Li, C.M., Weissman, J.S., Lipford, J.R., and Deshaies, R.J. (2022). Adaptive exchange sustains cullin-RING ubiquitin ligase networks and proper licensing of DNA replication. Proc Natl Acad Sci U S A 119, e2205608119. 10.1073/pnas.2205608119.

25. Cornish-Bowden, A. (1986). Why is uncompetitive inhibition so rare? A possible explanation, with implications for the design of drugs and pesticides. FEBS Lett 203, 3–6. 10.1016/0014-5793(86)81424-7.

26. Sato, Y., Yoshikawa, A., Yamagata, A., Mimura, H., Yamashita, M., Ookata, K., Nureki, O., Iwai, K., Komada, M., and Fukai, S. (2008). Structural basis for specific cleavage of Lys 63-linked polyubiquitin chains. Nature 455, 358–362. 10.1038/nature07254.

27. Suh, G.S., Poeck, B., Chouard, T., Oron, E., Segal, D., Chamovitz, D.A., and Zipursky, S.L. (2002). Drosophila JAB1/CSN5 acts in photoreceptor cells to induce glial cells. Neuron 33, 35–46. 10.1016/s0896-6273(01)00576-1.

28. Yu, C., Huszagh, A., Viner, R., Novitsky, E.J., Rychnovsky, S.D., and Huang, L. (2016). Developing a Multiplexed Quantitative Cross-Linking Mass Spectrometry Platform for Comparative Structural Analysis of Protein Complexes. Anal Chem 88, 10301–10308. 10.1021/acs.analchem.6b03148.

29. Gutierrez, C., Chemmama, I.E., Mao, H., Yu, C., Echeverria, I., Block, S.A., Rychnovsky, S.D., Zheng, N., Sali, A., and Huang, L. (2020). Structural dynamics of the human COP9 signalosome revealed by cross-linking mass spectrometry and integrative modeling. Proc Natl Acad Sci U S A 117, 4088–4098. 10.1073/pnas.1915542117.

30. Cao, S., Kang, S., Mao, H., Yao, J., Gu, L., and Zheng, N. (2022). Defining molecular glues with a dual-nanobody cannabidiol sensor. Nat Commun 13, 815. 10.1038/s41467-022-28507-1.

31. Wang, X., Chemmama, I.E., Yu, C., Huszagh, A., Xu, Y., Viner, R., Block, S.A., Cimermancic, P., Rychnovsky, S.D., Ye, Y., et al. (2017). The proteasome-interacting Ecm29 protein disassembles the 26S proteasome in response to oxidative stress. J Biol Chem 292, 16310–16320. 10.1074/jbc.M117.803619.

32. Fang, L., Kaake, R.M., Patel, V.R., Yang, Y., Baldi, P., and Huang, L. (2012). Mapping the protein interaction network of the human COP9 signalosome complex using a label-free QTAX strategy. Mol Cell Proteomics 11, 138–147. 10.1074/mcp.M111.016352.

33. Lv, L., Chen, P., Cao, L., Li, Y., Zeng, Z., Cui, Y., Wu, Q., Li, J., Wang, J.H., Dong, M.Q., et al. (2020). Discovery of a molecular glue promoting CDK12-DDB1 interaction to trigger cyclin K degradation. Elife 9. 10.7554/eLife.59994.

34. Słabicki, M., Kozicka, Z., Petzold, G., Li, Y.D., Manojkumar, M., Bunker, R.D., Donovan, K.A., Sievers, Q.L., Koeppel, J., Suchyta, D., et al. (2020). The CDK inhibitor CR8 acts as a molecular glue degrader that depletes cyclin K. Nature 585, 293–297. 10.1038/s41586-020-2374-x.

35. Khan, Z.M., Real, A.M., Marsiglia, W.M., Chow, A., Duffy, M.E., Yerabolu, J.R., Scopton, A.P., and Dar, A.C. (2020). Structural basis for the action of the drug trametinib at KSR-bound MEK. Nature 588, 509–514. 10.1038/s41586-020-2760-4.

36. Ryan, M.B., Quade, B., Schenk, N., Fang, Z., Zingg, M., Cohen, S.E., Swalm, B.M., Li, C., Özen, A., Ye, C., et al. (2024). The Pan-RAF-MEK Nondegrading Molecular Glue NST-628 Is a Potent and Brain-Penetrant Inhibitor of the RAS-MAPK Pathway with Activity across Diverse RAS- and RAF-Driven Cancers. Cancer Discov 14, 1190–1205. 10.1158/2159-8290.CD-24-0139.

37. Locke, J., Joseph, A.P., Peña, A., Möckel, M.M., Mayer, T.U., Topf, M., and Moores, C.A. (2017). Structural basis of human kinesin-8 function and inhibition. Proc Natl Acad Sci U S A 114, E9539–E9548. 10.1073/pnas.1712169114.

38. Tamayo, N.A., Bourbeau, M.P., Allen, J.R., Ashton, K.S., Chen, J.J., Kaller, M.R., Nguyen, T.T., Nishimura, N., Pettus, L.H., Walton, M., et al. (2022). Targeting the Mitotic Kinesin KIF18A in Chromosomally Unstable Cancers: Hit Optimization Toward an In Vivo Chemical Probe. J Med Chem 65, 4972–4990. 10.1021/acs.jmedchem.1c02030.

39. Shergalis, A.G., Marin, V.L., Rhee, D.Y., Senaweera, S., McCloud, R.L., Ronau, J.A., Hutchins, C.W., McLoughlin, S., Woller, K.R., Warder, S.E., et al. (2023). CRISPR Screen Reveals BRD2/4 Molecular Glue-like Degrader via Recruitment of DCAF16. ACS Chem Biol 18, 331–339. 10.1021/acschembio.2c00747.

40. Andersson, J., Cowland, S., Vestergaard, M., Yang, Y., Liu, S., Fang, X., Mukund, S., Ghimire-Rijal, S., Carter, C., Chung, G., et al. (2025). MTA-cooperative PRMT5 inhibitors from cofactor-directed DNA-encoded library screens. Proc Natl Acad Sci U S A 122, e2425052122. 10.1073/pnas.2425052122.

41. Mastronarde, D.N. (2005). Automated electron microscope tomography using robust prediction of specimen movements. J Struct Biol 152, 36–51. 10.1016/j.jsb.2005.07.007.

42. Wu, C., Huang, X., Cheng, J., Zhu, D., and Zhang, X. (2019). High-quality, high-throughput cryo-electron microscopy data collection via beam tilt and astigmatism-free beam-image shift. J Struct Biol 208, 107396. 10.1016/j.jsb.2019.09.013.

43. Punjani, A., Rubinstein, J.L., Fleet, D.J., and Brubaker, M.A. (2017). cryoSPARC: algorithms for rapid unsupervised cryo-EM structure determination. Nat Methods 14, 290–296. 10.1038/nmeth.4169.

44. Bepler, T., Morin, A., Rapp, M., Brasch, J., Shapiro, L., Noble, A.J., and Berger, B. (2019). Positive-unlabeled convolutional neural networks for particle picking in cryo-electron micrographs. Nat Methods 16, 1153–1160. 10.1038/s41592-019-0575-8.

45. Punjani, A., Zhang, H., and Fleet, D.J. (2020). Non-uniform refinement: adaptive regularization improves single-particle cryo-EM reconstruction. Nat Methods 17, 1214–1221. 10.1038/s41592-020-00990-8.

46. Punjani, A., and Fleet, D.J. (2021). 3D variability analysis: Resolving continuous flexibility and discrete heterogeneity from single particle cryo-EM. J Struct Biol 213, 107702. 10.1016/j.jsb.2021.107702.

47. Pettersen, E.F., Goddard, T.D., Huang, C.C., Couch, G.S., Greenblatt, D.M., Meng, E.C., and Ferrin, T.E. (2004). UCSF Chimera--a visualization system for exploratory research and analysis. J Comput Chem 25, 1605–1612. 10.1002/jcc.20084.

48. Emsley, P., Lohkamp, B., Scott, W.G., and Cowtan, K. (2010). Features and development of Coot. Acta Crystallogr D Biol Crystallogr 66, 486–501. 10.1107/S0907444910007493.

49. Jamali, K., Käll, L., Zhang, R., Brown, A., Kimanius, D., and Scheres, S.H.W. (2024). Automated model building and protein identification in cryo-EM maps. Nature 628, 450–457. 10.1038/s41586-024-07215-4.

50. Afonine, P.V., Poon, B.K., Read, R.J., Sobolev, O.V., Terwilliger, T.C., Urzhumtsev, A., and Adams, P.D. (2018). Real-space refinement in PHENIX for cryo-EM and crystallography. Acta Crystallogr D Struct Biol 74, 531–544. 10.1107/S2059798318006551.

51. Kao, A., Chiu, C.L., Vellucci, D., Yang, Y., Patel, V.R., Guan, S., Randall, A., Baldi, P., Rychnovsky, S.D., and Huang, L. (2011). Development of a novel cross-linking strategy for fast and accurate identification of cross-linked peptides of protein complexes. Mol Cell Proteomics 10, M110.002212. 10.1074/mcp.M110.002212.

52. Jiao, F., Yu, C., Wheat, A., Wang, X., Rychnovsky, S.D., and Huang, L. (2022). Two-Dimensional Fractionation Method for Proteome-Wide Cross-Linking Mass Spectrometry Analysis. Anal Chem 94, 4236–4242. 10.1021/acs.analchem.1c04485.

53. Tyanova, S., Temu, T., and Cox, J. (2016). The MaxQuant computational platform for mass spectrometry-based shotgun proteomics. Nat Protoc 11, 2301–2319. 10.1038/nprot.2016.136.

54. Wu, C., Ba, Q., Lu, D., Li, W., Salovska, B., Hou, P., Mueller, T., Rosenberger, G., Gao, E., Di, Y., et al. (2021). Global and Site-Specific Effect of Phosphorylation on Protein Turnover. Dev Cell 56, 111–124.e116. 10.1016/j.devcel.2020.10.025.

55. Demichev, V., Messner, C.B., Vernardis, S.I., Lilley, K.S., and Ralser, M. (2020). DIA-NN: neural networks and interference correction enable deep proteome coverage in high throughput. Nat Methods 17, 41–44. 10.1038/s41592-019-0638-x.

56. Kistner, F., Grossmann, J.L., Sinn, L.R., and Demichev, V. (2023). QuantUMS: uncertainty minimisation enables confident quantification in proteomics. bioRxiv, 2023.2006.2020.545604. 10.1101/2023.06.20.545604.

57. Pino, L.K., Searle, B.C., Bollinger, J.G., Nunn, B., MacLean, B., and MacCoss, M.J. (2020). The Skyline ecosystem: Informatics for quantitative mass spectrometry proteomics. Mass Spectrom Rev 39, 229–244. 10.1002/mas.21540.

